# EcDNA hubs drive cooperative intermolecular oncogene expression

**DOI:** 10.1101/2020.11.19.390278

**Authors:** King L. Hung, Kathryn E. Yost, Liangqi Xie, Sihan Wu, Joshua T. Lange, Connor V. Duffy, Katerina Kraft, Jun Tang, Quanming Shi, John C. Rose, M. Ryan Corces, Jeffrey M. Granja, Rui Li, Utkrisht Rajkumar, Robert Tjian, Vineet Bafna, Paul S. Mischel, Zhe Liu, Howard Y. Chang

**Affiliations:** Center for Personal Dynamic Regulomes, Stanford University School of Medicine, Stanford, CA 94305, USA; Janelia Research Campus, Howard Hughes Medical Institute, Ashburn, VA, USA; Department of Molecular and Cell Biology, Li Ka Shing Center for Biomedical and Health Sciences, CIRM Center of Excellence, University of California, Berkeley, CA, USA; Howard Hughes Medical Institute, Berkeley, CA, USA; Department of Cellular and Molecular Medicine, University of California at San Diego, La Jolla, CA, USA; Biomedical Sciences Graduate Program, University of California at San Diego, La Jolla, CA, 92093, USA; Department of Computer Science and Engineering, University of California at San Diego, La Jolla, CA, USA; Department of Pathology, University of California at San Diego, La Jolla, CA, USA; Howard Hughes Medical Institute, Stanford University School of Medicine, Stanford, CA 94305, USA

## Abstract

Extrachromosomal DNAs (ecDNAs) are prevalent in human cancers and mediate high oncogene expression through elevated copy number and altered gene regulation^1^. Gene expression typically involves distal enhancer DNA elements that contact and activate genes on the same chromosome^2,3^. Here we show that ecDNA hubs, comprised of ~10-100 ecDNAs clustered in the nucleus of interphase cells, drive intermolecular enhancer input for amplified oncogene expression. Single-molecule sequencing, single-cell multiome, and 3D enhancer connectome reveal subspecies of *MYC-PVT1* ecDNAs lacking enhancers that access intermolecular and ectopic enhancer-promoter interactions in ecDNA hubs. ecDNA hubs persist without transcription and are tethered by BET protein BRD4. BET inhibitor JQ1 disperses ecDNA hubs, preferentially inhibits ecDNA oncogene transcription, and kills ecDNA+ cancer cells. Two amplified oncogenes *MYC* and *FGFR2* intermix in ecDNA hubs, engage in intermolecular enhancer-promoter interactions, and transcription is uniformly sensitive to JQ1. Thus, ecDNA hubs are nuclear bodies of many ecDNAs tethered by proteins and platforms for cooperative transcription, leveraging the power of oncogene diversification and combinatorial DNA interactions. We suggest ecDNA hubs, rather than individual ecDNAs, as units of oncogene function, cooperative evolution, and new targets for cancer therapy.

## INTRODUCTION

Circular ecDNA encoding oncogenes is a prevalent feature of cancer genomes and potent driver of human cancer progression^4–7^. EcDNAs are covalently closed, double-stranded, and range from ~100 kilobases to several megabases in size^1,8–11^. EcDNAs lack centromeres and are randomly distributed among daughter cells after each cell division allowing for rapid accumulation and selection of ecDNA variants that drug resistance or other fitness advantage^5,12–14^. ecDNAs can evolve over time from submicroscopic episomes to large double minutes, and that these extrachromosomal elements can re-integrate into chromosomes and form tandem repeats termed homogeneously staining regions (HSRs)^15–19^. ecDNA possess increased chromatin accessibility and lack higher order chromatin compaction^1,20^, and encompass the endogenous oncogene enhancer elements^21^. ecDNA exists outside the normal chromosomal context by definition, and its spatial organization in the nucleus is poorly understood. Notably, ecDNAs can cluster during cell division or after DNA damage with unclear biological consequences^22–24^.

The *MYC* oncogene on human chromosome 8q24 is a hotspot for somatic DNA rearrangements in cancer^25^, and nearly 30% of *MYC* amplifications in human cancers exist as ecDNA^5^, typically encompassing *MYC* and 5’ portion of *PVT1* (plasmacytoma variant transcript 1)*. PVT1*, located 55kb 3’ of *MYC*, is a recurrent breakpoint in human cancers^26,27^. Structural rearrangements in *PVT1* often lead to the transcriptional activation of *MYC*, and historically led the way for the recognition of *MYC* as an oncogene^27,28^. The *PVT1* gene encodes a long noncoding RNA and contains multiple intragenic enhancers that normally interact with the *PVT1* promoter. However, when *PVT1* promoter is mutated or silenced, *PVT1* intragenic enhancers can activate *MYC* instead ^26^. The dynamic competition between *MYC* and *PVT1* promoters implicated *PVT1* promoter as a tumor suppressor DNA boundary element. Here we examine the spatial, epigenetic, and transcriptional dynamics of oncogenic ecDNAs, focusing on *MYC-PVT1* in human cancer cells, and reveal ecDNA hubs as combinatorial enhancer platforms for cooperative oncogene transcription.

## RESULTS

### ecDNA hubs are predominant sites of oncogene transcription

To understand the 3D context of ecDNA during transcription, we visualized the localization of ecDNAs in the nucleus during interphase using DNA fluorescence in situ hybridization (FISH)^29^ (**Figure 1a**). We designed probes targeting corresponding oncogenes and surrounding sequences amplified on ecDNAs in multiple ecDNA-containing cell lines including PC3 (*MYC*-amplified; prostate cancer), COLO320-DM (*MYC*-amplified; colorectal carcinoma), HK359 (*EGFR*-amplified; glioblastoma multiforme), and SNU16 (*MYC*- and *FGFR2*-amplified; gastric cancer)^1^ (**Supplemental Figure 1a**). To validate that these sequences are located on ecDNA molecules, we performed DNA FISH on metaphase spreads to confirm the presence of tens to hundreds individual extrachromosomal DNAs per cell based on DNA FISH signal located outside of metaphase chromosomes (**Supplemental Figure 1b**). In a subset of cell lines, we also employed two-color DNA FISH to interrogate a neighboring control locus *in cis* that is not amplified on ecDNA (**Supplemental Figure 1a**); chromosomal copies of the oncogene appear as paired dots with neighboring chromosomal loci proximal to each other (**Figure 1a**) while ecDNAs have a single color. Two-color FISH experiment in an ecDNA-negative cell line, HCC1569, consistently showed paired two-color signals of similar sizes as expected from the chromosomal loci (**Supplemental Figure 1c**). In contrast, in all ecDNA-positive cancer cells we assessed, DNA FISH signal for ecDNAs was largely restricted to specific areas of the nucleus of interphase cells despite arising from tens to hundreds of individual ecDNA molecules, suggesting that ecDNAs strongly clustered with one another, a feature we term ecDNA hubs (**Figure 1a**). These ecDNA hubs occupy a much larger space than neighboring chromosomal segments of the same size, suggesting that they are composed of many ecDNA molecules clustering tightly in space. We used the pair autocorrelation function g(r) (Methods) to measure the spatial distribution of ecDNA hubs. g(r) estimates the probability of detecting another ecDNA signal at increasing distances from the viewpoint of an index ecDNA signal and is equal to 1 for a uniform, random distribution. This quantification showed a significant increase in ecDNA clustering over short distances (0-40 pixels, 0-1.95 microns, **Figure 1b**), with all cell lines and oncogenes displaying increased autocorrelation compared to a simulated random distribution (Methods). EcDNA clusters were much larger than diffraction limited spots (~0.3 microns), consistent with co-localization of multiple ecDNA copies within individual clusters. To confirm these findings in live cells, we generated a Tet-operator (TetO) array knock-in of the *MYC* ecDNA in COLO320-DM cells, and labeled ecDNA with TetR-eGFP (TetO-eGFP COLO320-DM) (**Figure 1c**, Methods). Live cell imaging revealed multiple bright and locally accumulated signal in the nucleus likely corresponding to ecDNA hubs composed of clustered ecDNA molecules, whereas the TetR-eGFP signal is homogeneously distributed in the parental cell line without the TetO array (**Figure 1c**, **Supplemental Figure 1d, Supplemental Movie 1**). EcDNA hubs are dynamic in living cells; they both move and change shape with time. We note that TetR-eGFP labeled ecDNA hubs are spatially more compact in living cells than in DNA FISH studies of fixed cells, likely due to the fact that the TetO array is not integrated in all ecDNA molecules, as well as potential differences induced by denaturation during DNA FISH. These results suggest that ecDNA clustering occurs across various cancer types with different oncogene amplifications.

**Figure 1.**
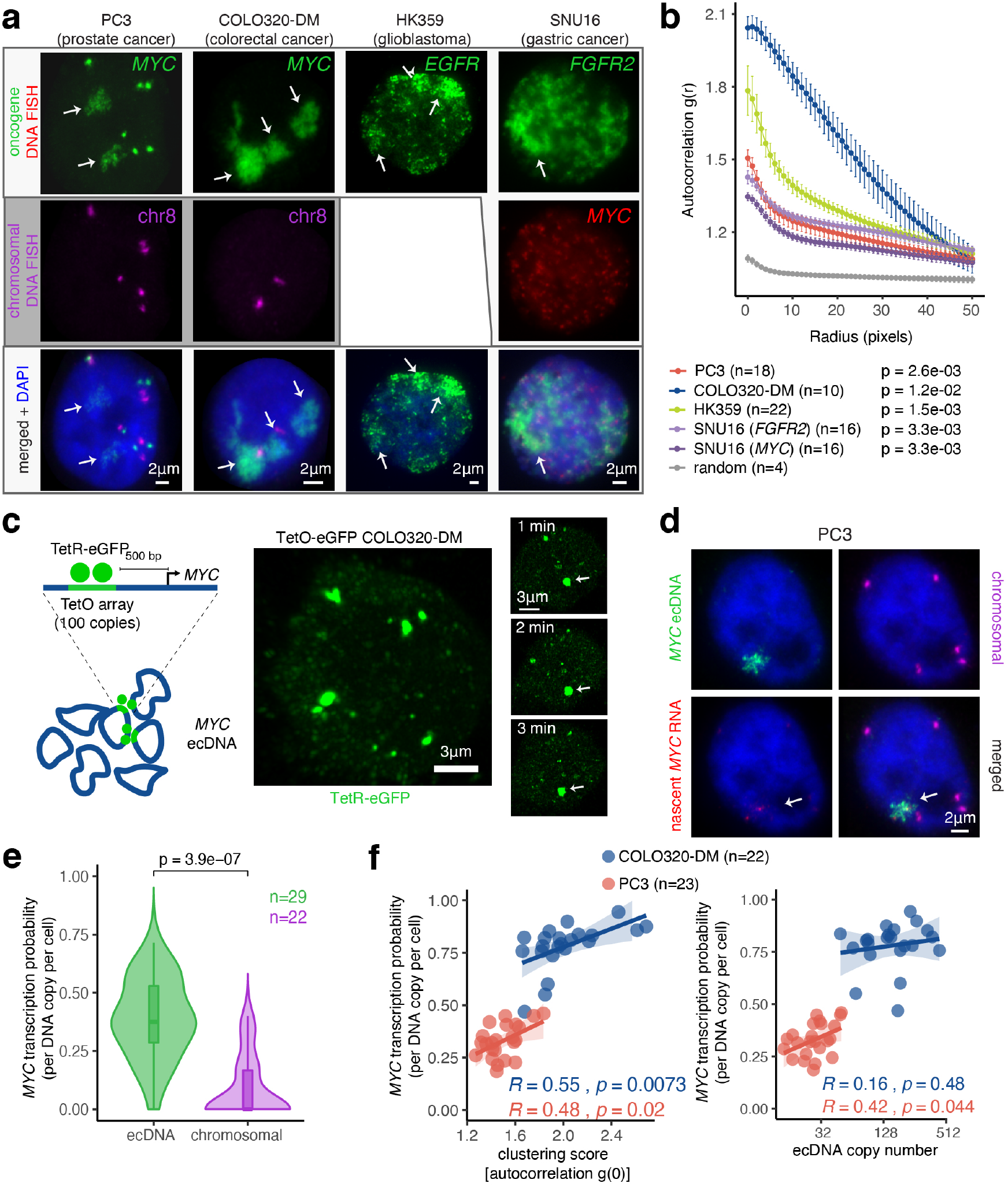
EcDNA imaging reveals ecDNA clustering and cooperative transcriptional bursting. **(a)** Representative FISH images showing ecDNA clustering during interphase in PC3 (*MYC* ecDNA, 1.5 Mb FISH probe), COLO320-DM (*MYC* ecDNA, 1.5 Mb FISH probe), HK359 (*EGFR* ecDNA) and SNU16 (*MYC* and *FGFR2* ecDNA, 200 kb *MYC* FISH probe) cells. For PC3 and COLO320-DM cells, a FISH probe targeting an adjacent chromosomal locus was also used. Scale bars, 2 μm. **(b)** Quantification of ecDNA clustering in interphase cells by autocorrelation (g) at different pixel radii (r). Data are represented as mean ± SEM at each radial value. A p-value comparing each cell line with random distribution is listed. P-values were calculated using the Wilcoxon test comparing the autocorrelation function values at radius = 0. **(c)** Schematic for ecDNA imaging based on TetO array knock-in and labeling with TetR-eGFP (left). Representative images of TetR-eGFP signal in TetO-eGFP COLO320-DM cells at indicated timepoint during live cell imaging time course (right). Scale bars, 3 μm. **(d)** Representative image from combined DNA FISH for *MYC* ecDNA (100 kb probe) and adjacent chromosomal DNA with nascent *MYC* RNA FISH in PC3 cells. Scale bars, 2 μm. **(e)** Quantification of *MYC* transcription probability measured by nascent RNA FISH in (D) normalized to DNA copy number measured by DNA FISH for ecDNA and chromosomal loci (box center line, median; box limits, upper and lower quartiles; box whiskers, 1.58x interquartile range, here and throughout). **(f)** Scatter plots of ecDNA copy number (measured by DNA FISH) or ecDNA clustering score (measured by autocorrelation of ecDNA FISH signal at r = 0) versus *MYC* transcription probability measured by nascent RNA FISH. Correlation coefficients calculated using Pearson’s R.

To assess the transcriptional activity of ecDNA hubs, we combined DNA and nascent RNA FISH in the PC3 and COLO320-DM cell lines to visualize actively transcribing *MYC* alleles as colocalized DNA and RNA FISH signal, using a short 100 kb Oligopaint DNA FISH probe to enable quantification of individual ecDNA molecules within ecDNA hubs (**Figure 1d, Supplemental Figure 1a,e–f**, Methods). We quantified colocalized RNA and DNA FISH signals to compute the probability of *MYC* transcription from each ecDNA molecule in a snapshot of time (Methods); this measure accounts for the copy number of ecDNA template and subsumes the frequency and duration of *MYC* transcription. In the PC3 cell line, we were able to more confidently distinguish chromosomal *MYC* transcription from ecDNA-derived transcription due to lower ecDNA copy numbers (**Figure 1d**). Interestingly, the majority of nascent *MYC* mRNA transcripts colocalized with ecDNA clusters rather than the chromosomal locus and transcription probability is significantly higher from ecDNA relative to the chromosomal locus (**Figure 1d,e**). Quantification of nascent *MYC* transcription from ecDNAs vs ecDNA clustering (using the autocorrelation function of ecDNA FISH signal, Methods) showed a significant correlation between *MYC* transcription probability and ecDNA clustering (*R*=0.48-0.55, *p*< 0.05 for both PC3 and COLO320-DM, **Figure 1f** left). EcDNA clustering is a better predictor of ecDNA transcription probability than ecDNA copy number (n=14-50 in PC3, n=49-441 in COLO320-DM, *R*=0.16-0.42, **Figure 1f** right). Thus, each ecDNA molecule is more likely to transcribe the oncogene when more ecDNA copies are in the same cell, especially in the form of ecDNA hubs.

### Single-cell co-variation identifies ecDNA enhancers associated with potent oncogene expression

To understand regulation of oncogene expression on ecDNAs, we set out to identify regulatory elements on ecDNAs that correlate with high oncogene expression. While previous data suggest that ecDNAs contribute to cancer cell heterogeneity^6,9^, the chromatin regulatory landscape of ecDNAs and its relationship to oncogene transcription has not been studied on a single-cell level. To address this, we focused on a pair of colorectal cancer cell lines, COLO320-DM and COLO320-HSR, which were derived from the same patient tumor and therefore contain highly similar genetic backgrounds except for the context in which *MYC* is amplified^17^. *MYC* is amplified on ecDNA in COLO320-DM cells versus tandem chromosomal amplicons (HSRs) in COLO320-HSR cells (**Supplemental Figure 2a**). Given the heterogeneous nature of ecDNAs across COLO320-DM cells, we hypothesized that we could exploit the cell-to-cell variation in ecDNA sequence to identify regulatory elements whose activity positively predicts potent ecDNA oncogene expression in individual cells. We took a single-cell multiomic approach and performed a droplet-based, paired assay for transposase-accessible chromatin using sequencing (ATAC-seq) and RNA sequencing in the same single cell (**Supplemental Figure 2b**,**c**, Methods) and obtained paired transcriptomic and chromatin accessibility profiles from a total of 72,049 cells. Cells were first visualized with uniform manifold approximation and projection (UMAP)^30^ independently based on either transcriptomic or chromatin accessibility profiles (Methods). UMAPs of either single-cell ATAC-seq or single-cell RNA-seq data showed separate clustering of COLO320-DM and COLO320-HSR cell lines as expected (**Figure 2a**). We then integrated the transcriptomic and chromatin accessibility profile for each cell to interrogate how chromatin accessibility covaries with gene expression. For each cell, we calculated a gene accessibility score for *MYC*, which incorporates ATAC-seq signals from the gene body and those from distal regulatory elements^31^ (Methods). Accessibility scores for *MYC* increased with RNA expression (**Supplemental Figure 2d**; Pearson R = 0.25, p < 2.2e-16). RNA expression as well as accessibility scores for *MYC* were highly heterogeneous in the ecDNA *MYC*-amplified COLO320-DM cell population relative to the chromosomal HSR *MYC*-amplified COLO320-HSR population (**Figure 2b**). These observations suggest that variable activities of regulatory elements may explain cell-to-cell variation in oncogene expression by ecDNAs.

**Figure 2.**
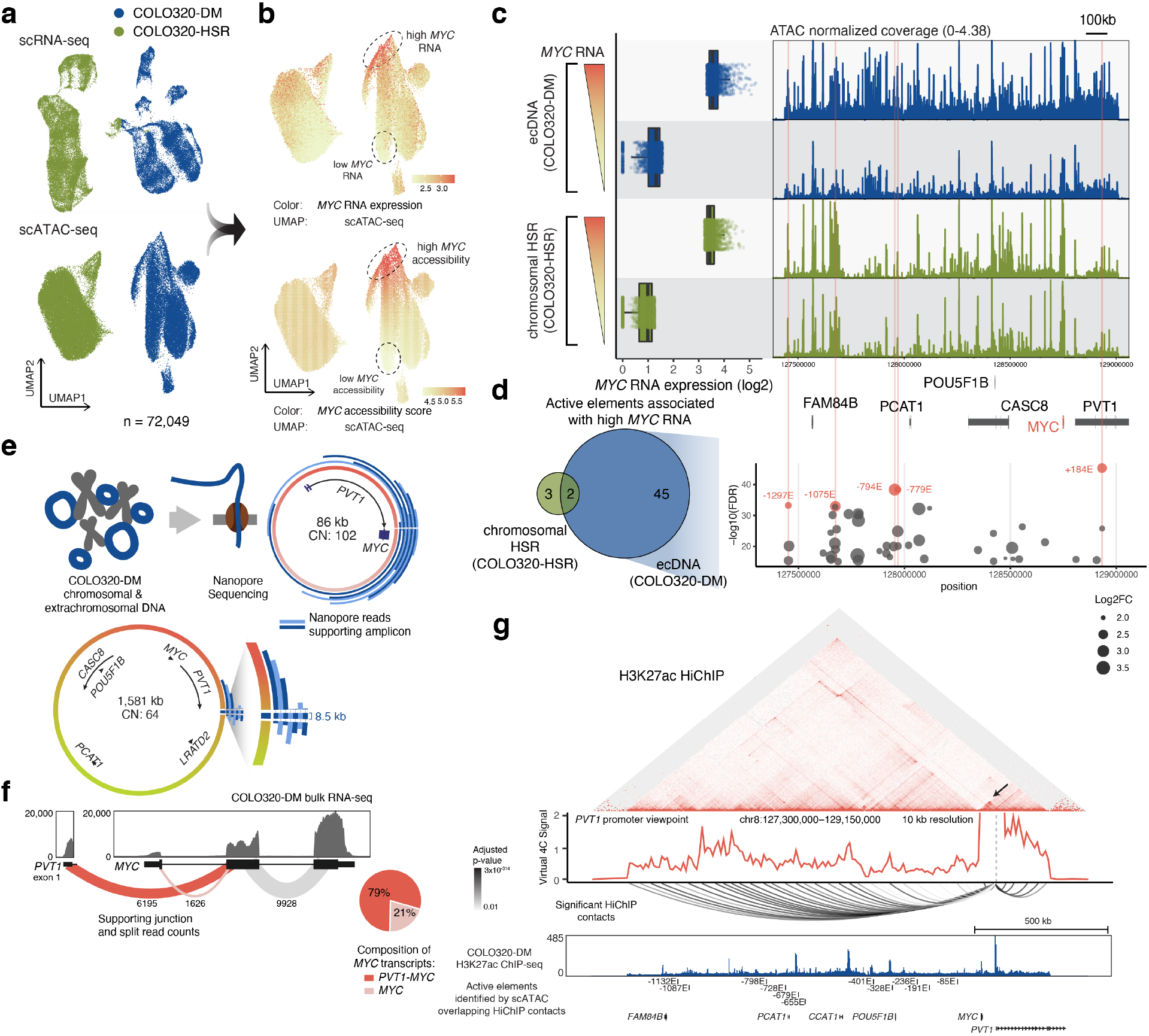
Genomic dissection identifies ecDNA enhancers associated with high *MYC* expression, ecDNA sequence heterogeneity and enhancer-promoter contacts. **(a)** COLO320-DM and COLO320-HSR cells profiled using single-cell paired RNA and ATAC-seq visualized with UMAP based on either the RNA or the ATAC-seq data. **(b)** Normalized *MYC* RNA expression levels were visualized on the ATAC-seq UMAP (top). An accessibility score for *MYC* was calculated using the ATAC-seq data and visualized on the UMAP with a color scale (bottom). **(c)** Cells were binned based on *MYC* RNA expression as outlined in Supplemental Figure 2b, and *MYC* expression levels of the top and bottom bins of COLO320-DM and COLO320-HSR are shown in a boxplot (left). The normalized ATAC-seq coverage for each bin is shown in the same order (right). **(d)** A venn diagram on the left shows number of variable elements identified on ecDNA amplicons in COLO320-DM compared to that identified on chromosomal HSRs in COLO320-HSR. 45 variable elements were uniquely seen on ecDNA and 2 variable elements overlap with chromosomal HSRs. All 47 variable elements on ecDNA were plotted across the entire amplified region as a dot plot, in which the x-axis represents genomic coordinates, the y-axis shows −log10 of the false discovery rate (FDR), and the size of each dot represents the log2 fold change (right). Five of these elements labelled and colored in red are the most significantly variable elements. They were named based on their relative position in kilobases in reference to the *MYC* transcriptional start site (TSS), with a negative number denoting an element 5’ of MYC and a positive number denoting an element 3’ of *MYC*. The regions occupied by the marked elements were also highlighted in the coverage tracks correspondingly. **(e)** Schematic for nanopore sequencing of COLO320-DM cells (top left). Nanopore reads supporting *MYC* containing amplicon structures shown with alternating colors. **(f)** Bulk RNA sequencing coverage from COLO320-DM with exon-exon junction spanning read counts shown (top). Pie chart quantifying relative abundance of full-length *MYC* and fusion *PVT1-MYC* transcripts using the read count supporting either junction (bottom). **(g)** H3K27ac HiChIP contact matrix (5 kb resolution, square root coverage normalized) in COLO320-DM cells centered on *MYC* locus (top). Arrow designates increased HiChIP signal at 86 kb *PVT1-MYC* fusion amplicon. Virtual 4C plots from *PVT1* promoter viewpoint with significant chromatin contacts identified by FitHiChIP shown below (middle). ATAC-seq signal track with active regulatory elements identified by scATAC-seq associated with high *MYC* expression that overlap loop anchors highlighted (bottom). Active elements named based on their relative position in kilobases in reference to the *MYC* transcriptional start site (TSS) as in panel c.

To identify active regulatory elements on ecDNAs in cells that express high levels of *MYC*, we assigned cells into 20 bins based on *MYC* RNA levels and adjusted ATAC-seq signals from each bin using copy numbers calculated from background coverage to perform pairwise testing of differential ATAC-seq peaks between the top and the bottom RNA expression bin (**Supplemental Figure 2b**, Methods). A comparison of ATAC-seq coverage tracks suggested that high *MYC*-expressing COLO320-DM cells contain both higher ecDNA copy number based on increased background signal as well as a number of increasingly accessible elements, while chromosomal HSRs displayed similar chromatin accessibility profiles in the top and bottom RNA bins of COLO320-HSR cells (**Figure 2c**). Differential peak analysis identified 47 active elements on ecDNA that were strongly associated with high *MYC* expression compared to 5 active elements on chromosomal HSRs (**Figure 2d**). These findings suggest that increased *MYC* transcription is associated with increased chromatin accessibility at numerous distal regulatory elements in the context of ecDNA. The most significantly upregulated DNA elements (named by their distance to the *MYC* promoter) demonstrated accessibility that scaled with *MYC* mRNA expression when accounting for DNA copy number only in the ecDNA context (**Supplemental Figure 2e**, zoomed in). Notably, significantly active elements were distributed throughout the entire amplified region. Most of these active elements were concentrated in the interval between *FAM84B* and *CASC8*, consistent with increased accessibility of regulatory elements 5’ of the *MYC* coding sequence across colon adenocarcinomas^32^. As expected, cells with high *MYC* expression tend to have higher ecDNA copy numbers; however, we note that there is a high level of variability in ecDNA copy number for cells with similar levels of *MYC* RNA output, suggesting that copy number alone does not fully predict RNA expression (**Supplemental Figure 2f,g**). Importantly, the decreased accessibility at regulatory elements in the low RNA bin of COLO320-DM is not likely to be caused by differences in data quality, as we did not observe a decrease in overall TSS enrichment (**Supplemental Figure 2h**). Moreover, accessibility profiles of high and low *MYC* RNA bins in COLO320-HSR cells are highly similar, demonstrating that the strong co-variation of regulatory elements with *MYC* expression and their heterogeneous activities within COLO320-DM cells may be a unique feature of ecDNA (**Figure 2c,d**). Together, these results indicate that differential enhancer usage may be key to understanding cellular heterogeneity in ecDNA oncogene expression.

### Combinatorial intermolecular contacts between enhancers and promoters drive ecDNA diversification and oncogene expression

With the discoveries that (i) ecDNA hubs in the nucleus are associated with active oncogene expression and (ii) oncogene expression by ecDNAs is linked to differential regulatory elements, we next interrogated whether these differential elements participate in novel enhancer-promoter interactions among potentially distinct ecDNA molecules. While one may predict that each ecDNA molecule must contain all of the sequence elements (enhancers, promoters, etc.) to promote oncogene expression *in cis*, the recognition of ecDNA hubs where multiple ecDNA molecules come into spatial proximity raises the possibility that DNA regulatory elements may cooperate among several ecDNAs to enable oncogene expression. In this second scenario, it may be possible to observe selection for ecDNA structures which lack canonical *cis* regulatory elements necessary for oncogene expression as well as ecDNA structures without coding oncogene elements if these distinct molecules can cooperate within transcriptionally active hubs. Here we integrate whole genome sequence analysis, single-molecule DNA sequencing, and 3D enhancer connectome mapping to distinguish between these two scenarios.

To reconstruct ecDNA molecules in COLO320-DM cells, we obtained whole-genome sequencing data and applied AmpliconArchitect, a computational tool which uses both copy number variation and structural variant analysis of short sequencing reads to reconstruct DNA amplicons arising from ecDNAs and other complex rearrangements^5,6,33^. We identified multiple “subspecies” of ecDNA amplicons resulting from the rearrangement of the *MYC* locus in COLO320-DM cells with a range of copy numbers (**Supplemental Figure 3a,b**). Notably, we detected abundant ecDNAs which lacked either the *MYC* coding sequence and contained only regulatory DNA elements, or conversely, contained a truncated *MYC* coding sequence and lacked distal regulatory elements (**Supplemental Figure 3b**). Next, we used nanopore-based single-molecule sequencing to obtain long contiguous ecDNA reads (mean 9 kb, maximum 201 kb; **Supplemental Figure 3c**). We observed high concordance between structural variants detected by long and short-read sequencing (**Supplemental Figure 3d**) and confirmed that one of the most abundant rearrangements of the *MYC* locus generates an 86kb amplicon that comprises the promoter and exon 1 of noncoding RNA *PVT1* fused to exons 2 and 3 of *MYC* (**Figure 2e**). Here the *PVT1* promoter replaces the *MYC* promoter to generate a *PVT1-MYC* fusion transcript and a new 5’ UTR for *MYC*, which is hypothesized to overcome *PVT1* and *MYC* promoter competition^26^. Previous studies have demonstrated that a functional protein isoform of MYC can initiate from a start codon in exon 2, suggesting that transcripts derived from the fusion *PVT1-MYC* transcript generate functional MYC protein^34,35^. However, the amplicon containing the *PVT1-MYC* fusion in COLO320-DM cells is not predicted to be covalently linked to many of the active regulatory elements located 5’ of the *MYC* coding sequence. We also confirmed multiple junctions present in a predicted ~1.58Mb amplicon with full-length *MYC* coding sequence that retains many distal regulatory elements but is predicted to be present at a lower copy number (64 instead of 102 for the 86kb amplicon). We observed multiple single-molecule sequencing reads spanning an 8.5 kb segment predicted to join two opposite ends of the larger amplicon segment (**Figure 2e**).

Our results suggest a high degree of heterogeneity among ecDNA molecules in COLO320-DM, with two major subspecies possessing unique variants of the *MYC* coding sequence as well as distinct connectivity to putative regulatory elements. Previous studies have also noted the existence of the *PVT1-MYC* fusion in COLO320-DM cells, originally as a 5’ structural abnormality of the *MYC* gene observed by Northern blot analysis^36–38^ and more recently with sequencing based methods^39^. We confirmed by RNA-seq analysis that the majority (79%) of the *MYC* mRNA transcripts in COLO320-DM cells arise from the *PVT1-MYC* fusion present on the 86 kb amplicon (predicted to be 61% of *MYC* ecDNA by DNA copy number) (**Figure 2f**). Single-cell transcriptomic co-variation analysis also validated *PVT1-MYC* transcription in COLO320-DM; reads mapping to *PVT1* positively correlated with reads mapping to *MYC* in COLO320-DM cells but not COLO320-HSR cells (Pearson R = 0.996 for COLO320-DM, R = - 0.909 for COLO320-HSR; **Supplemental Figure 3e**). Thus, in a cancer cell line harboring a variety of ecDNA species, oncogene output appears to be preferentially driven by a specific ecDNA subspecies. Moreover, the rearrangement that creates the 86 kb amplicon excludes known *MYC* enhancers both 5’ of the *MYC* transcriptional start site or 3’ in the *PVT1* locus, raising the question of whether and how ecDNA-encoded *PVT1-MYC* accesses distal enhancers.

Next, we mapped active enhancers and their target genes in COLO320-DM cells using HiChIP, a protein-directed 3D genome conformation assay^40,41^, targeting histone H3 lysine 27 acetylation (H3K27ac) associated with active enhancers and enriched on ecDNA as observed by immunofluorescence in metaphase cells^1^. These results first provided independent confirmation of the 86 kb *PVT1-MYC* amplicon, which is evident as a tightly interacting core on the HiChIP map with increased vertex signal following correction for copy number variation with square root coverage normalization (**Figure 2g**, arrow, Methods). Second, we identified a number of active enhancer elements identified by single-cell ATAC-seq originally located 5’ of the *MYC* gene with significant contact to the *PVT1*/*PVT1-MYC* promoter (**Figure 2g, Supplemental Figure 3f**). We noted that while the canonical *MYC* promoter participates in several focal enhancer contacts (**Supplemental Figure 3g**), HiChIP signal at the *PVT1* promoter is elevated across the entirety of the amplified region, supported by a uniform density of highly significant chromatin contacts (**Figure 2g**). While several loops overlap structural rearrangements identified by long-read sequencing, many high-confidence loops identified by HiChIP are independent of structural arrangements, suggesting they represent true enhancer-promoter contacts (**Supplemental Figure 3d**). Collectively, these results suggest an extensive degree of ecDNA primary sequence diversification, including structural variants that would be predicted to limit contact *in cis* with distal regulatory elements. However, robust transcription associated with combinatorial and intermolecular enhancers usage in within ecDNA hubs led us to speculate that multiple species of ecDNA molecules may cooperate when in spatial proximity to achieve robust transcription of cargo genes.

### BRD4 bridges ecDNA hubs and drives hub oncogene transcription

The extensive long-range and H3K27ac-associated DNA contacts raised the possibility that BET proteins may be involved in transcription from ecDNA hubs. Bromodomain and extraterminal domain (BET) proteins are chromatin reader proteins that recognize H3K27ac and are intricately involved in enhancer function. Genomic regions with multiple contiguous enhancers, termed super enhancers or stretch enhancers, are recognized based on their extensive decoration by H3K27ac, transcription coactivator complex Mediator, and BET proteins such as BRD4^42,43^. The *MYC* gene is flanked by tissue-specific super enhancers in certain cancers and BRD4 occupancy as well as *MYC* transcription are highly sensitive to the BET inhibitor JQ1^44^, which displaces BET proteins from H3K27ac^45^. We also previously showed that BRD4 occupancy marked the winner in promoter competition between *PVT1* and *MYC*^26^. To determine the role of BET proteins in ecDNA-derived transcription, we examined BRD4 occupancy at the *MYC* locus in both COLO320-DM and COLO320-HSR cells. Endogenous epitope tagging of BRD4 combined with ecDNA labeling with TetR-eGFP showed that BRD4 colocalized with ecDNAs in COLO320-DM cells (**Figure 3a**). This colocalization of GFP and BRD4 signals was not observed in cells without TetO integration (**Supplemental Figure 4a)**. Chromatin immunoprecipitation and sequencing (ChIP-seq) of H3K27ac, BRD4, and ATAC-seq in COLO320-DM and COLO320-HSR cells showed that indeed H3K27ac peaks, marking active ecDNA enhancers, are also occupied by BRD4, including many active elements associated with high *MYC* transcription and in contact with the *PVT1-MYC* promoter (−1132E, −1087E, −679E, −655E, −401E, −328E, −85E) (**Supplemental Figure 4b**). Importantly, the *PVT1* promoter is one of the highest sites of BRD4 occupancy in ecDNA containing cells (COLO320-DM) but not in cells with chromosomal amplification (COLO320-HSR) (**Figure 3b, Supplemental Figure 4c**). Whole genome sequencing showed that HSR amplicons in COLO320-HSR do not contain the *PVT1* promoter, as evidenced by reduced whole genome sequencing coverage (**Figure 3b**, WGS track). Thus, ecDNAs and chromosomal amplicons in this cell line pair have adopted two diametrically opposite strategies to inactivate the tumor suppressor *PVT1* promoter – cooption of *PVT1* promoter to drive MYC transcription in ecDNA versus deletion of *PVT1* promoter in chromosomal *MYC* amplicons.

**Figure 3.**
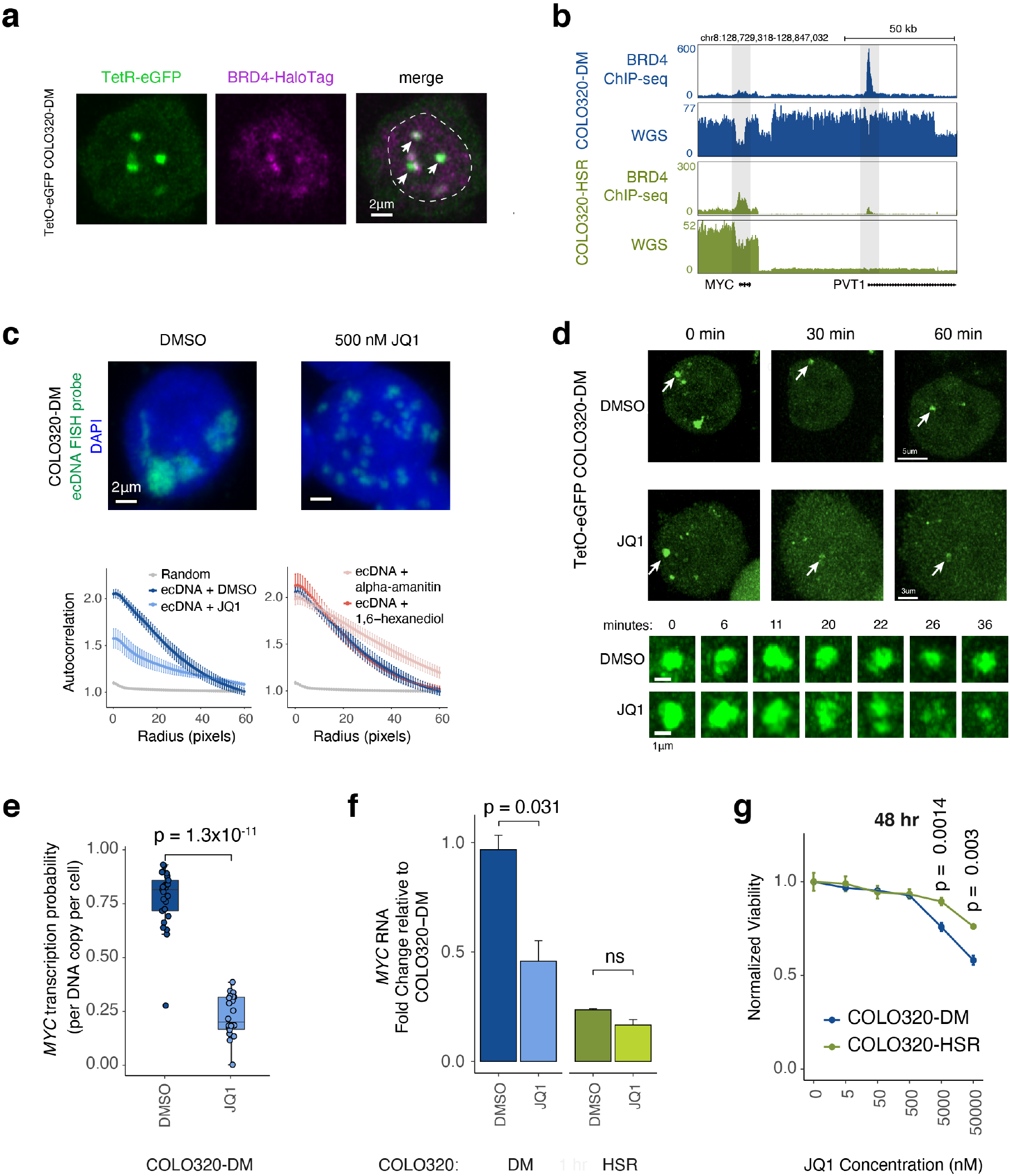
BET proteins mediate ecDNA hub formation and transcription. **(a)** Representative image of eDNA labeled with TetR-eGFP in TetO-eGFP COLO320-DM cells overlaid with BRD4-HaloTag signal. Dashed line indicates nucleus boundary. **(b)** BRD4 ChIP-seq and whole genome sequencing coverage at *MYC* and *PVT1* loci in COLO320-DM and COLO320-HSR cells. **(c)** Representative DNA FISH images for *MYC* ecDNA in interphase COLO320-DM cells treated with either DMSO or 500 nM JQ1 for 6 hours (top). Quantification of ecDNA clustering in interphase cells by autocorrelation (g) at different pixel radii (r) for COLO320-DM cells treated either with DMSO, 500 nM JQ1, 1% 1,6-hexanediol, or 100 μg/mL alpha-amanitin for 6 hours. Error bars represent standard error between individual cells (n = 10) used for quantification (bottom). **(d)** Representative images of TetR-eGFP signal in TetO-eGFP COLO320-DM cells treated either with DMSO or 500 nM JQ1 at indicated timepoint during live cell imaging time course. (**e)** Quantification of *MYC* transcription probability measured by nascent RNA FISH normalized to ecDNA copy number measured by DNA FISH for COLO320-DM cells treated either with DMSO or 500 nM JQ1 for 6 hours (n = 22-25). **(f)** Quantification of *MYC* RNA measured by RT-qPCR for COLO320-DM and COLO320-HSR cells treated either with DMSO or 500 nM JQ1 for 6 hours (n = 2). **(g)** Normalized cell viability for COLO320-DM and COLO320-HSR cells treated with a range of JQ1 concentrations for 48 hours normalized to cell viability for DMSO treated cells (n = 3).

Next, we evaluated the consequences of BET inhibitor JQ1 on ecDNA hubs, the dominant site of *MYC* transcription in COLO320-DM. Treatment with 500 nM JQ1 dramatically dispersed ecDNA hubs in COLO320-DM after 6 hours (**Figure 3c**); large ecDNA hubs split into multiple small ecDNA signals within the nucleoplasm, as reflected in reduced autocorrelation g(r) across multiple length scales (**Figure 3c**). However, ecDNA dispersal by JQ1 appears to be a highly specific effect. Transcription inhibition by either the RNA polymerase II inhibitor alpha-amanitin or treatment with 1,6-hexanediol, a hydrophobic compound that can dissolve certain nuclear condensates^46^, both of which robustly reduce MYC transcription (**Supplemental Figure 5a,b**), did not affect ecDNA hubs (**Figure 3c**, **Supplemental Figure 5c**). Live cell imaging with TetO-GFP COLO320-DM cells revealed that JQ1 causes ecDNA hubs to disperse after ~30 minutes of drug treatment while ecDNA hubs in control cells remain intact (**Figure 3d, Supplemental Movie 2,3**). Together, these results suggest that bromodomain-H3K27ac interaction of BET proteins is essential for ecDNA clustering.

We next examined the effect of JQ1 on ecDNA-derived oncogene transcription. JQ1 treatment strongly reduced *MYC* transcription probability by four-fold per ecDNA copy, as evidenced by joint nascent RNA and DNA FISH (**Figure 3e**, **Supplemental Figure 5a**). Because BET proteins are also involved in *MYC* transcription from chromosomal DNA, we further compared the effect of JQ1 on COLO320-DM vs COLO320-HSR cells. BRD4 ChIP-seq showed that JQ1 treatment equivalently dislodged BRD4 genome-wide in these isogenic cells (**Supplemental Figure 5d**). Nonetheless, treatment with 500 nM JQ1 preferentially lowered *MYC* mRNA level in COLO320-DM cells, a dose which had no significant effect on *MYC* mRNA level in COLO320-HSR cells (**Figure 3f**). Dose titration of JQ1 confirmed the preferential killing of COLO320-DM cells over HSR cells (**Figure 3g**). These results demonstrate a unique dependence on BET proteins for ecDNA hub formation; disruption of ecDNA hubs cause preferential suppression of *MYC* oncogene transcription and death of ecDNA-bearing cancer cells.

In contrast to the ability of BETi to inhibit ecDNA transcription, we found that ecDNAs appear resistant to targeting of individual enhancers. We and others have previously used CRISPR interference (CRISPRi) with targeted catalytically dead Cas9 fused to KRAB silencer domain to identify functional *MYC* enhancers in the chromosomal context ^26,41,47^. CRISPRi of the *PVT1* promoter, but not *MYC* promoter, indeed reduced *PVT1-MYC* transcription and total *MYC* mRNA level in COLO320-DM, confirming the promoter cooption from the 86kb ecDNA subspecies and serving as positive control (**Supplemental Figure 5e**). In contrast, individually targeting 6 enhancers with high BRD4 occupancy on ecDNA did not significantly reduce bulk *MYC* mRNA levels (**Supplemental Figure 5e,f**). These results suggest that ecDNA dispersal may be a more effective strategy to overcome highly cooperative oncogene enhancers in ecDNA hubs.

### EcDNA hubs support intermolecular cross regulation of two oncogene loci

As several lines of evidence presented above suggest that ecDNA hubs may promote intermolecular enhancer-promoter interactions, we investigated whether these interactions can be precisely mapped and perturbed. Due to the overlap of DNA segments that compose ecDNA amplicons in the COLO320-DM cell line, uniquely mapping enhancer elements to distinct molecules is challenging. To overcome this, we focused on a human gastric cancer cell line, SNU16, which contains two major types of ecDNAs. One type of ecDNA contains a *MYC* amplicon derived from chromosome 8 and the other contains an *FGFR2* amplicon derived from chromosome 10. Image analysis of dual-color metaphase FISH in SNU16 cells with EcSeg showed that these two amplicons are located on distinct molecules in the majority of ecDNAs (~35% of all ecDNA molecules contain *MYC* only, and ~60% contain *FGFR2* only; **Figure 4a,b**)^48^, which was also supported by analysis of whole-genome sequencing by AmpliconArchitect (**Supplemental Figure 6a,b**). In contrast, *MYC* and *FGFR2* ecDNAs intermingle with each other during interphase in hubs as demonstrated by increased colocalization in dual-color interphase FISH (**Figure 4c,d**). These data led us to hypothesize that ecDNA hubs in SNU16 may allow intermolecular interactions between *MYC*-bearing and *FGFR2*-bearing ecDNAs. To test this idea, we performed H3K27ac HiChIP on SNU16 cells to map enhancer-promoter interactions. Because a small percentage of ecDNA molecules appeared to contain both *MYC* and *FGFR2* (~5%, **Figure 4b**), we used AmpliconArchitect to identify regions that were predicted to be fused on ecDNA, where we observe a high density of HiChIP signal potentially resulting from *cis* interactions (**Figure 4e**, dashed box highlighting translocation). Outside of this region, we identified intermolecular contacts between active enhancers from the *MYC* super enhancer to *FGFR2* promoter, as well as enhancer contacts from *FGFR2* amplicon back to *MYC* (**Figure 4e**, arrows). These intermolecular interactions remain prominent following correction for copy number variation with square root coverage normalization.

**Figure 4.**
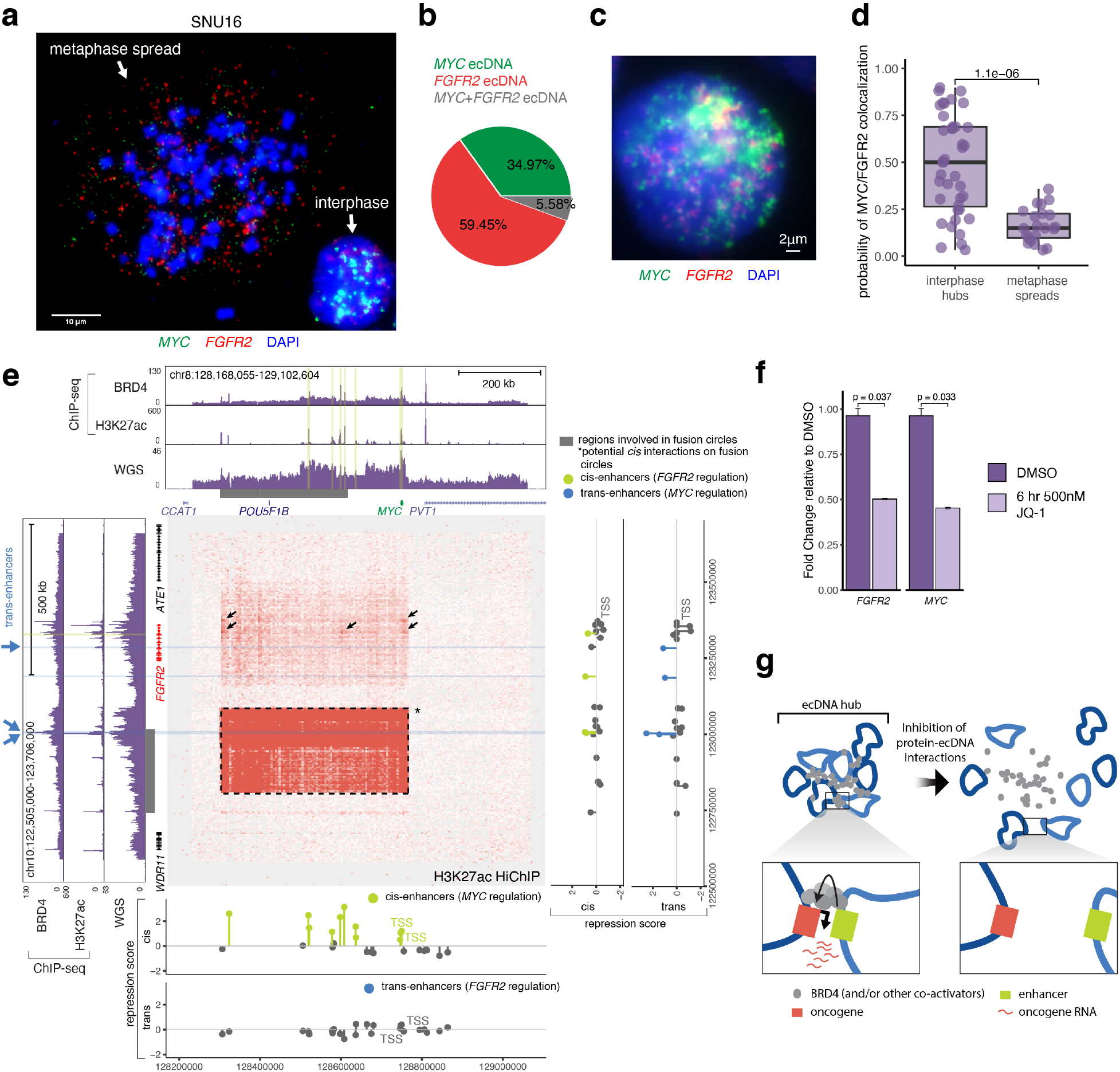
Intermolecular interaction among ecDNA molecules. **(a)** Representative FISH images showing extrachromosomal *MYC* and *FGFR2* amplifications in metaphase spreads in SNU16 cells. **(b)** Quantification of *MYC*, *FGFR2*, and dual oncogene ecDNAs in SNU16 cells detected by DNA FISH in metaphase cells. **(c)** Representative DNA FISH images showing clustering of *MYC* and *FGFR2* ecDNA during interphase in SNU16 cells. Scale bars, 2 μm. **(d)** Quantification of *MYC* and *FGFR2* colocalization observed by DNA FISH in interphase and metaphase SNU16 cells. **(e)** H3K27ac HiChIP contact matrix (5 kb resolution, square root coverage normalized) in SNU16 cells between *MYC* and *FGFR2* loci with 1D H3K27ac ChIP-seq, BRD4 ChIP-seq, and whole genome sequencing coverage tracks shown. Regions predicted to be covalently linked based on AmpliconArchitect analysis annotated in grey and region of potential *cis* interaction indicated by asterisk and dashed outline on HiChIP contact matrix. Area with focal contacts between *MYC* and *FGFR2* loci highlighted. Lollipop plots (bottom and right) show the repressive effects of CRISPR interference on *MYC* or *FGFR2* expression along the genomic track. Guides targeting putative enhancers that had a repressive effect on the oncogene in the same locus are marked as cis-enhancer targets (light green), whereas guides that repressed the oncogene in the other locus are marked as trans-enhancer targets (blue). Details of how the repression score was calculated can be found in Methods and Supplemental Figure 6c. Plots show data from two biological replicates. **(f)** Quantification of *MYC* and *FGFR2* RNA measured by RT-qPCR for SNU16 cells treated either with DMSO or 500 nM JQ1 for 6 hours. **(g)** A schematic diagram of the proposed ecDNA hub model for intermolecular cooperation. EcDNA hubs bring many heterogeneous ecDNA molecules into close proximity, allowing for interactions between enhancers and oncogenes on distinct molecules and promoting oncogene RNA transcription. The formation of these ecDNA hubs is mediated by BRD4 and/or other proteins that likely bind strongly to the endogenous sequences that are amplified on ecDNAs. Inhibition of protein-ecDNA interactions leads to dispersal of ecDNAs, which in turn reduces oncogene transcription.

To assess the effects of perturbing intermolecular enhancer-promoter interactions on each respective oncogene, we stably expressed dCas9-KRAB in SNU16 cells (SNU16-dCas9-KRAB) and delivered lentiviral pools of sgRNA guides targeting regulatory elements located on either *MYC*-bearing ecDNAs or *FGFR2*-bearing ecDNAs (**Supplemental Figure 6c**). After guide infection and antibiotic selection, cells were labeled with *MYC* or *FGFR2* mRNA targeting FISH probes, signal was amplified with PrimeFlow, and RNA expression of each oncogene was analyzed by flow cytometry. We sorted on cell fractions that had negative, low or high expression of either oncogene, extracted genomic DNA and performed targeted sequencing of the lentiviral guide pools (**Supplemental Figure 6c**). Based on guide abundances in each cell fraction compared to unsorted cells, we calculated a combined repression score that summarizes the degree to which each guide is enriched in cells with low oncogene expression and depleted in cells with high oncogene expression (Methods, **Supplemental Figure 6c**). A high repression score would suggest that the targeted enhancer normally upregulates oncogene expression. Using this approach, we found that CRISPRi of a subset of H3K27ac- and BRD4-bound enhancer regions 5’ of *MYC* as well as *MYC* TSS inhibited *MYC* RNA expression, while CRISPRi of loci not bound by H3K27ac or BRD4 had no effect on expression (**Figure 4e**, bottom panel, **Supplemental Figure 6d**). CRISPRi of enhancers on MYC ecDNAs had minimal effect on *FGFR2* expression. Conversely, CRISPRi of 5’ and intragenic enhancers of *FGFR2* led to decreases in both *FGFR2 in cis* and *MYC* expression *in trans* (**Figure 4e**, right panel, **Supplemental Figure 6d**), suggesting that enhancers derived from the *FGFR2* locus may activate transcription from *MYC* ecDNAs. Given that a small percentage of ecDNAs contain sequences from both the *MYC* and the *FGFR2* loci (**Figure 4b,e**), two of the intergenic *FGFR2* enhancers that affect *MYC* transcription lie in the predicted translocated region, and thus could be either *cis*- or *trans*-acting. However, the *FGFR2* intragenic enhancer is required for *MYC* expression and is located outside of the predicted fusion circles and is a bona-fide *trans*-acting enhancer (**Figure 4e**). This *FGFR2* ecDNA enhancer is also located within a broad region with increased intermolecular interactions with *MYC* ecDNAs as shown by increased HiChIP signals. Finally, treatment with the BET protein inhibitor JQ1 led to concomitant reduction in both *MYC* and *FGFR2* expression (**Figure 4f**). These observations suggest that colocalization of *MYC*- and *FGFR2*-encoding ecDNAs in hubs may be important for the expression of both oncogenes.

## DISCUSSION

In this study, we show that oncogene-carrying ecDNAs in cancer cells strongly colocalize in clusters that associate with high levels of transcription, a phenomenon we term ecDNA hubs. This local congregation of ecDNAs promotes novel enhancer-promoter interactions and oncogene expression (**Figure 4g**). In turn, variable usage of these enhancers across molecules strongly contributes to cell-to-cell heterogeneity in oncogene-driven programs. Unlike chromosomal transcription hubs which are restricted to local *cis* regulatory elements, chromatin conformation data and CRISPR interference suggest that ecDNA hubs may also involve *trans* regulatory elements arising on distinct ecDNA molecules. This discovery has profound implications in 1) how ecDNAs undergo selection and 2) how rewiring of oncogene regulation on ecDNA contributes to oncogenic transcription and cancer cell heterogeneity.

### EcDNA hubs in oncogene selection and cancer cell heterogeneity

EcDNA molecules are products of stringent genetic selection which are able to drive high levels of oncogene expression outside of the normal chromosomal context and provide a fitness advantage to cancer cells that harbor them. Given that ecDNA has been separated from the 3D genomic context of its chromosomal origin, it has been proposed that the co-selection of oncogenes and enhancers shapes ecDNA amplicon structures^21^. In this study, we presented evidence for intermolecular interactions among ecDNA molecules carrying distinct enhancer elements. With this new observation, we propose a two-level model for oncogene-enhancer co-selection. The first level of co-selection occurs to individual ecDNAs; molecules that possess functional enhancers can promote oncogene expression and provide better fitness to cancer cells compared to ones that do not. The second level of co-selection occurs to the repertoire of ecDNAs in hubs. In other words, we predict that individual ecDNA molecules are not required to contain all of the enhancers necessary for promoting oncogene expression; rather, they exist as part of an ecDNA hub that facilitates chromatin interactions among a diverse repertoire of regulatory elements and promotes interactions between the target oncogene and functional enhancers which may be located on distinct molecules. This model raises the intriguing concept that winning the clonal *competition* among cancer cells occurs through clonal *cooperation* among ecDNA molecules. This type of evolutionary dynamics has been documented in viruses, where cooperation of a mixture of specialized variants outperforms a pure population of wild type virus^49,50^. Furthermore, our ecDNA cooperation model predicts that mutations on individual molecules may be better tolerated if functional elements are present on other molecules in a hub. If true, this tolerance may increase ecDNA sequence diversity and permit rare mutations, including those that confer resistance to therapies. Others have previously reported ecDNA mutational diversity and rapid response to environmental changes^51^, though further investigation is needed to measure mutational diversity in functional enhancers on ecDNAs.

Our study shows that enhancer usage can be highly variable on ecDNAs, which associates with cancer cell heterogeneity in oncogene activity. This may be attributed to differential enhancer-promoter interactions which occur in the context of ecDNA hubs. As dozens of ecDNA molecules can cluster together in many possible spatial configurations, oncogene promoters may have a greater opportunity to “sample” various enhancers via novel enhancer-promoter interactions. When different ecDNAs arise from two different chromosomes, such as in SNU16 cells, they intermingle in ecDNA hubs that enable ectopic enhancer-promoter interactions among oncogene loci that normally do not occur on linear chromosomes. We speculate that these differential interactions contribute to the highly variable enhancer activities and enhancer rewiring on ecDNAs.

### *PVT1* and *MYC* regulation

*PVT1,* specifically its promoter element, emerges as a strong selective pressure that shapes ecDNA and HSR amplicons. Most copies of the *MYC-PVT1* amplicon in COLO320-HSR cells have focal deletions of the *PVT1* promoter, suggesting that a *PVT1* promoter deletion was either an early mutation or provided a strong selective advantage. This observation is consistent with experimental data that a focal *PVT1* promoter deletion, as small as six bases, can activate *MYC* expression in the chromosomal context and confer a growth advantage^26^. A different strategy is adopted in COLO320-DM cells, in which a rearrangement fuses the *PVT1* promoter to the *MYC* oncogene; the fusion *PVT1-MYC* transcript becomes the dominant *MYC* mRNA in COLO320-DM cells. These appear as two divergent strategies for overcoming promoter competition between *PVT1* and *MYC* and amplifying *MYC* expression.

### Potential mechanisms of ecDNA hub formation

What may be the forces that drive ecDNA hub formation? Transcriptional hubs are thought to be nuclear condensates formed by high concentration of transcription factors, mediator, and RNA polymerase II through interaction of low complexity protein sequences and RNA^52–54^. Small, transient hubs of several hundred protein molecules are distinguished from stable condensates formed by liquid-liquid phase separation that are 10-20 times larger and restrict the movement of molecules into and out of the condensate (reviewed by ^55^). Small transient transcriptional hubs are necessary for gene transcription^56^, and we speculate that ecDNA hubs are a kind of transcriptional hub mediated by protein-protein interaction involving BRD4 and likely additional proteins. *MYC* ecDNA hubs are not disrupted by transcriptional inhibition with alpha-amanitin nor by 1,6-hexanediol, suggesting that ecDNA hubs do not depend on RNA polymerase II or specific IDR-IDR interactions sensitive to hexanediol such as Mediator 1^46^. BRD4 bromodomains have also been report to mediate recruitment into transcription hubs independent of IDR-IDR interactions^57,58^, and *BRD4-NUT* translocation and overexpression in rare human cancers can cause interchromosomal interactions^59^. Interestingly, McSwiggen *et al.* have previously proposed a nuclear compartmentalization model for Herpes Simplex Virus replication compartments in which accessible viral DNA outcompetes host chromatin for the transcriptional machinery and creates high local concentrations of RNA polymerase II^60^. We speculate that ecDNA hubs may exploit via a similar mechanism, wherein accessible ecDNAs sponge up BRD4 and DNA-binding proteins locally to create this uneven spatial distribution.

BET proteins are required for ecDNA hub maintenance and oncogene transcription in the COLO320-DM and SNU16 models. As BET proteins can normally concentrate accessible DNA, exclude heterochromatin, and mediate long-range enhancer-promoter communication^58,61^, our results suggest that ecDNA hubs may coopt endogenous mechanisms of long-range gene looping *within* chromosomes to promote *intermolecular* chromatin interactions in ecDNA ensembles. While we have focused on *MYC* in this study, our model predicts that other ecDNA oncogenes may also exploit their endogenous enhancer mechanisms to operate in ecDNA hubs. As others have shown that functional enhancers are co-selected with *EGFR* on ecDNAs in glioblastoma^21^, we speculate that proteins that mediate endogenous enhancer-EGFR interactions could be involved in ecDNA hub maintenance as well.

### Implications for cancer cell evolution and therapeutic opportunities

Our results suggest that ecDNA hubs provide a palatable explanation for well-known tendency of a subset of ecDNA+ cancer cells to develop HSRs. A recent study has shown that double-strand breaks in ecDNAs can trigger aggregation, micronucleus formation, and reintegration into chromosomal HSRs^24^. Rather than independently suffering concurrent DNA breaks and integrating into the same chromosomal locus, ecDNAs that are spatially proximal in hubs could enable correlated DNA breaks^62^ and concentrated DNA cargo, creating a potential set up for HSR formation. Finally, ecDNA hubs may impact ecDNA replication and segregation. Previous work has demonstrated that ecDNAs are transmitted into daughter cells in clusters during mitosis^23^. Interestingly, papillomavirus episomes hitchhike into daughter cells during mitosis by tethering to segregating chromosomes via BRD4^63^. Future studies may address whether an ecDNA hub serves as a unit of inheritance or merely as a transient congregation. The observation of ecDNA hubs also warrants further investigation into whether this unusual 3D organization of DNA molecules impacts other cellular processes regulated by genome organization, such as DNA repair and replication.

The recognition that ecDNA hubs promote oncogene transcription may provide new therapeutic opportunities. While chromosomal DNA amplicons are covalently linked on an HSR, ecDNA hubs are held together by proteins. In the case of the colorectal COLO320-DM cell line, we show that BET protein inhibition by JQ1 disaggregates ecDNA hubs and reduces ecDNA-derived *MYC* expression. JQ1 preferentially inhibited *MYC* transcription from ecDNA+ cancer cells and inhibited both *MYC* and *FGFR2* in SNU16 cells with dual oncogene hubs. Given that ecDNA hubs are associated with high transcription, the observation that ecDNA hub maintenance depends on BRD4 and/or other DNA-binding proteins may present a unique vulnerability of ecDNA-driven cancers. The specific protein that mediates ecDNA clustering may be cancer-type specific, as it likely needs to bind strongly to the endogenous sequences of the ecDNA-amplified regions. Future studies using a broad screening approach coupled with analysis of ecDNA hub maintenance may identify proteins that mediate of ecDNA clustering in various cancer types and will be highly informative for potential therapeutic efforts.

---

**Supplemental Figure 1.**
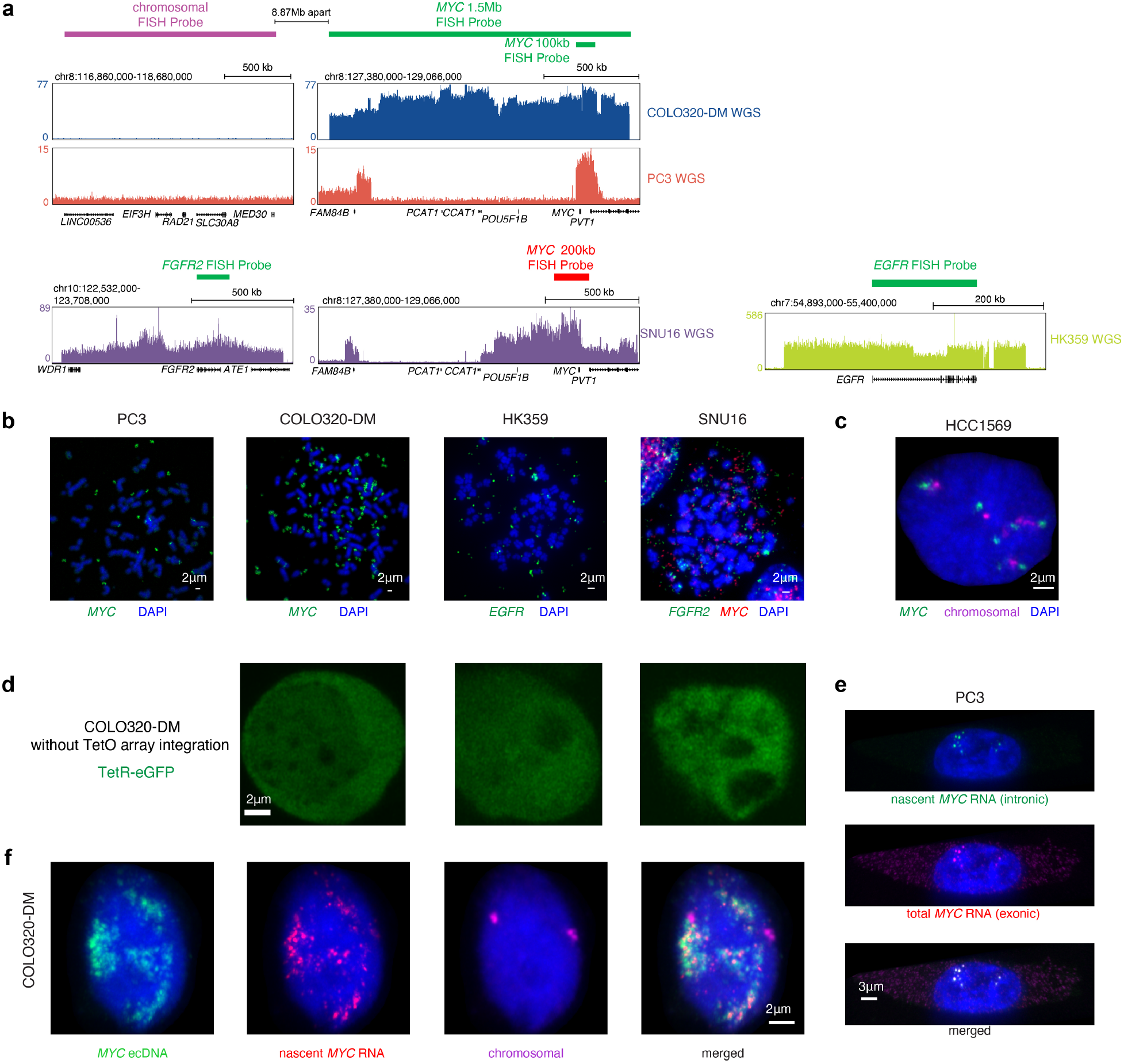
EcDNA FISH probe design and validation of extrachromosomal FISH signal. **(a)** Whole genome sequencing signal tracks for PC3, COLO320-DM, HK359 and SNU16 cells with DNA FISH probe locations highlighted. For analysis of COLO320-DM and PC3 three FISH probes were used: a 1.5 Mb Oligopaint FISH probe tiling the entire amplified locus surrounding *MYC* in COLO320-DM cells which was used for clustering analysis in Figure 1a,b; a 100 kb Oligopaint FISH probe at the *MYC* coding sequence used for ecDNA copy number quantification in Figure 1d,e,f; and a 1.5 Mb Oligopaint FISH probe tiling an unamplified region located 8.87 Mb upstream of the *MYC* locus on chromosome 8. For DNA FISH in SNU16 and HK359 cells, pre-designed FISH probes targeting either *EGFR*, *FGFR2* or *MYC* were used. **(b)** Representative FISH images showing extrachromosomal amplifications in metaphase spreads for (*MYC* ecDNA, 200 kb probe), COLO320-DM (*MYC* ecDNA, 200 kb probe), HK359 (*EGFR* ecDNA) and SNU16 (*MYC* and *FGFR2* ecDNA, 200 kb *MYC* probe) cells. Scale bars, 2 μm. **(c)** Representative DNA FISH images using chromosomal and 1.5 Mb *MYC* probes in non-ecDNA amplified cell line HCC1569 during interphase. Scale bars, 2 μm. **(d)** Representative images of TetR-eGFP signal in COLO320-DM cells without TetO array integration. Scale bars, 2 μm. **(e)** Representative images of nascent *MYC* RNA FISH probe validation showing overlap of nascent *MYC* RNA (intronic) FISH probe and total *MYC* RNA (exonic) FISH probe in PC3 cells. Scale bars, 3 μm. **(f)** Representative images from combined DNA FISH for *MYC* ecDNA (100 kb probe) and adjacent chromosomal DNA with nascent *MYC* RNA FISH in COLO320-DM cells. Scale bars, 2 μm.

**Supplemental Figure 2.**
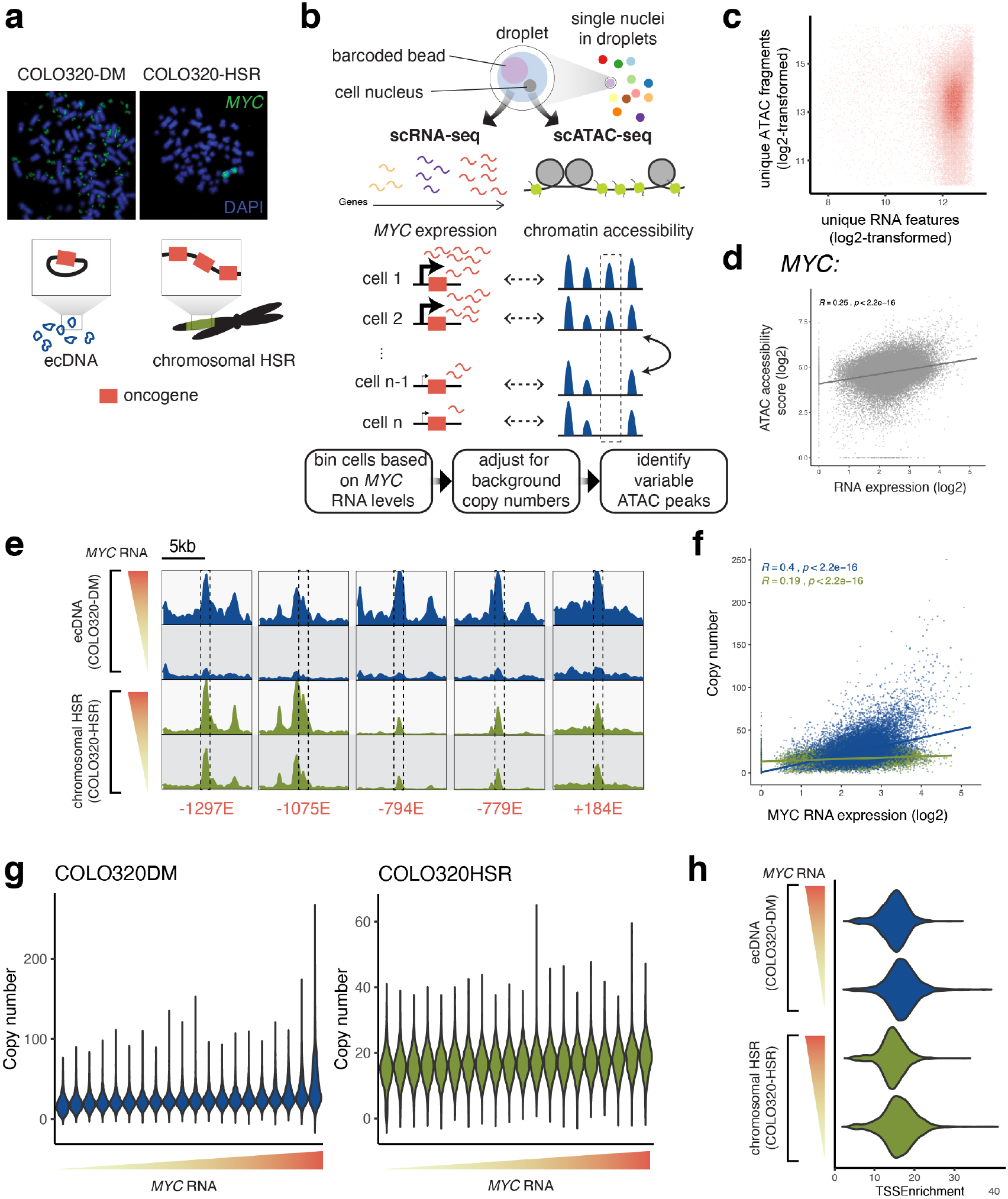
Schematic and validation of paired single-cell RNA-seq and ATAC-seq analysis in COLO320-DM and COLO320-HSR. **(a)** Representative metaphase FISH images showing extrachromosomal *MYC* signals in COLO320-DM and chromosomal tandem *MYC* signals in COLO320-HSR. The schematic at the bottom illustrates oncogene amplification on circular ecDNA in COLO320-DM and chromosomal HSRs in COLO320-HSR. **(b)** Schematic showing the single-cell assay design and analysis. Droplets containing single nuclei and barcoded beads were generated on the 10X Genomics platform, and RNA and ATAC-seq reads were obtained from each single cell to simultaneously assay gene expression and chromatin accessibility. Cells were binned based on MYC RNA levels, the copy number of ecDNA or HSR amplicons for each cell was adjusted based on background ATAC signals, and variable ATAC-seq peaks were identified by differential marker testing between the top and the bottom bin. **(c)** Cells passing ATAC-seq and RNA-seq quality control filters are plotted based on unique ATAC-seq fragments and unique RNA features (both log2-transformed). **(d)** An accessibility score for *MYC* was calculated from the ATAC-seq data and plotted against normalized *MYC* expression from RNA-seq data, showing a positive correlation. **(e)** Zoom-ins of the ATAC-seq coverage of each of the five most significantly variable elements identified in Figure 2d. Dashed boxes mark the positions of these elements. **(f)** A scatter plot of estimated *MYC* amplicon copy numbers and normalized log2-transformed *MYC* expression of all individual cells showing positive correlations and a high level of copy number variability. **(g)** Estimated *MYC* amplicon copy number distributions of all RNA bins of COLO320-DM and COLO320-HSR. **(h)** A violin plot showing distributions of TSS enrichments in the high and low RNA bins of COLO320-DM and COLO320-HSR.

**Supplemental Figure 3.**
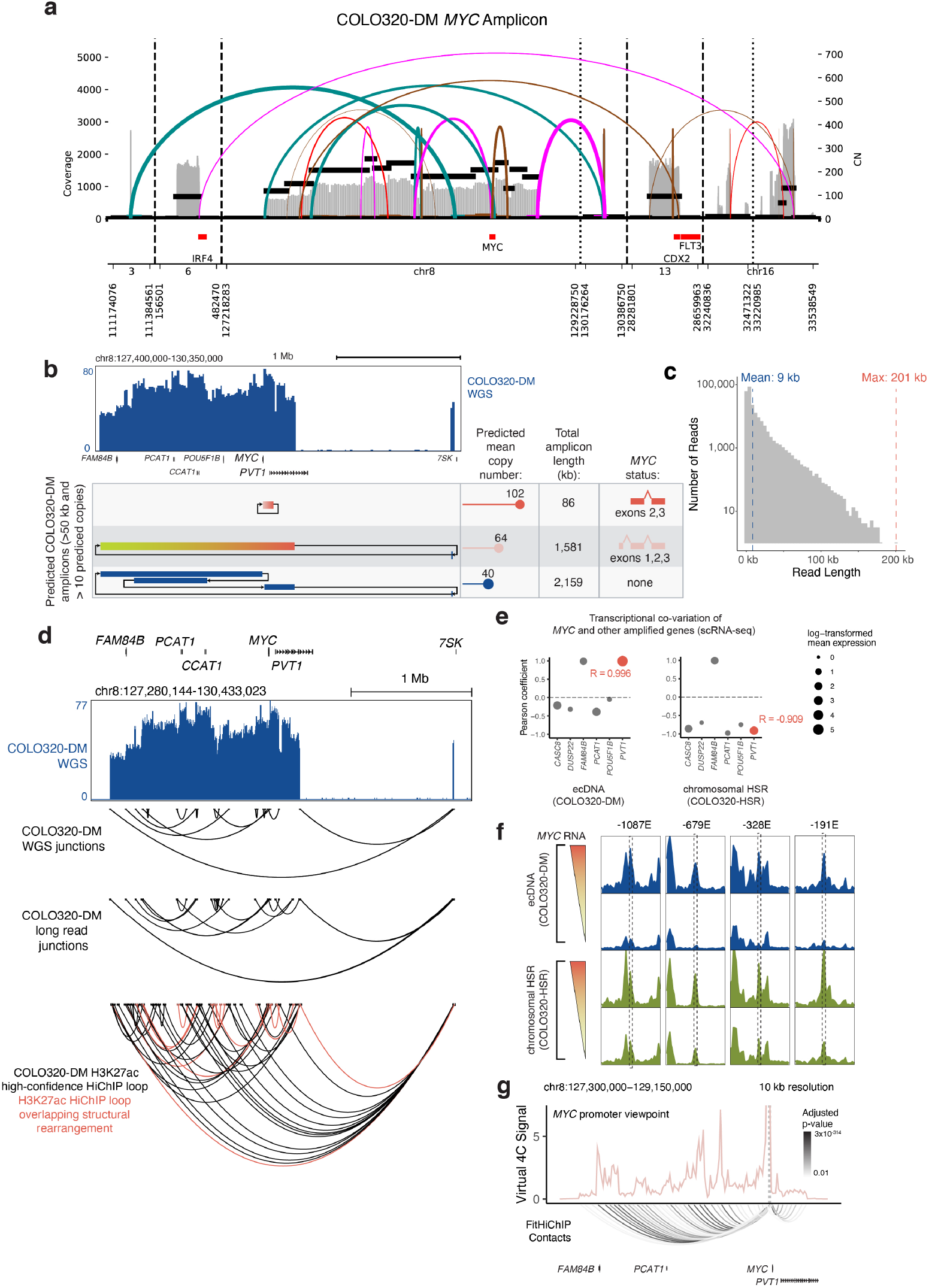
ecDNA sequence heterogeneity and enhancer-promoter contacts. **(a)** Structural variant (SV) view of AmpliconArchitect (AA) reconstruction of the *MYC* amplicon in COLO320-DM cells. **(b)** Whole genome sequencing coverage from COLO320-DM and amplicon structures predicted by AmpliconArchitect, annotated with predicted mean copy number, total amplicon length, and *MYC* status. DNA segments shown as blocks with arrows designating junctions supported by whole genome sequencing. Amplicon structures with over 50 kb in total length, over 10 predicted copies, and with head-to-tail orientation characteristic of circular amplicons shown. **(c)** Distribution of read lengths from long-read nanopore sequencing of COLO320-DM. **(d)** Whole genome sequencing coverage from COLO320-DM (top). Junctions detected by whole genome sequencing, junctions detected by long-read nanopore sequencing, and H3K27ac HiChIP high-confidence loops identified by HICCUPS shown below, with HiChIP loops overlapping structural rearrangements highlighted. **(e)** Pearson correlation of transcription co-variation from scRNA-seq quantification of *MYC* and other amplified genes in COLO320-DM and COLO320-HSR cells. **(f)** Aggregate scATAC-seq signal tracks from top *MYC* RNA expression bin and bottom *MYC* RNA expression bin in COLO320-DM and COLO320-HSR cells, with differential peaks associated with high *MYC* RNA expression and overlapping HiChIP loop anchors highlighted. **(g)** COLO320-DM H3K27ac HiChIP virtual 4C plots from the *MYC* promoter viewpoint with FitHiChIP loops shown below, colored by adjusted p-value.

**Supplemental Figure 4.**
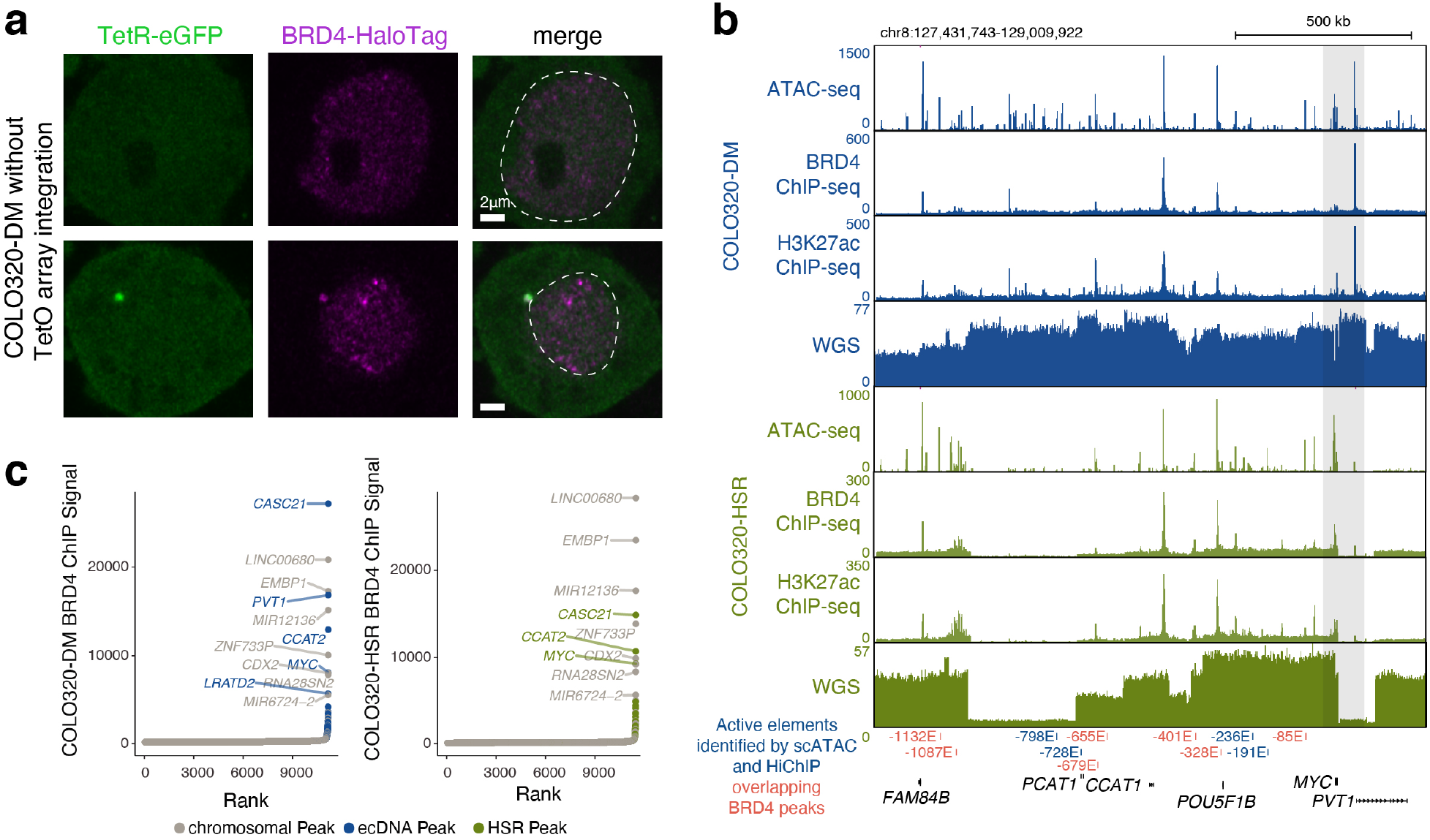
ecDNA is enriched for BRD4 occupancy. **(a)** Representative image of TetR-eGFP signal in COLO320-DM cells without TetO array integration overlaid with BRD4-HaloTag signal. Dashed line indicates nucleus boundary. We noted cytoplasmic TetR-eGFP signal in a subset of COLO320-DM cells without TetO array integration during co-detection of BRD4-HaloTag (bottom) but these signals were located outside of the nuclear boundary and did not colocalize with BRD4-HaloTag. **(b)** ATAC-seq, BRD4 ChIP-seq, H3K27ac ChIP-seq and whole genome sequencing coverage tracks for COLO320-DM and COLO320-HSR cells at amplified MYC locus. Active regulatory elements identified in Figure 3D shown below with elements overlapping BRD4 ChIP-seq peaks highlighted. **(c)** Ranked BRD4 ChIP-seq signal for COLO320-DM and COLO320-HSR, which peaks included in either ecDNA or HSR amplifications highlighted.

**Supplemental Figure 5.**
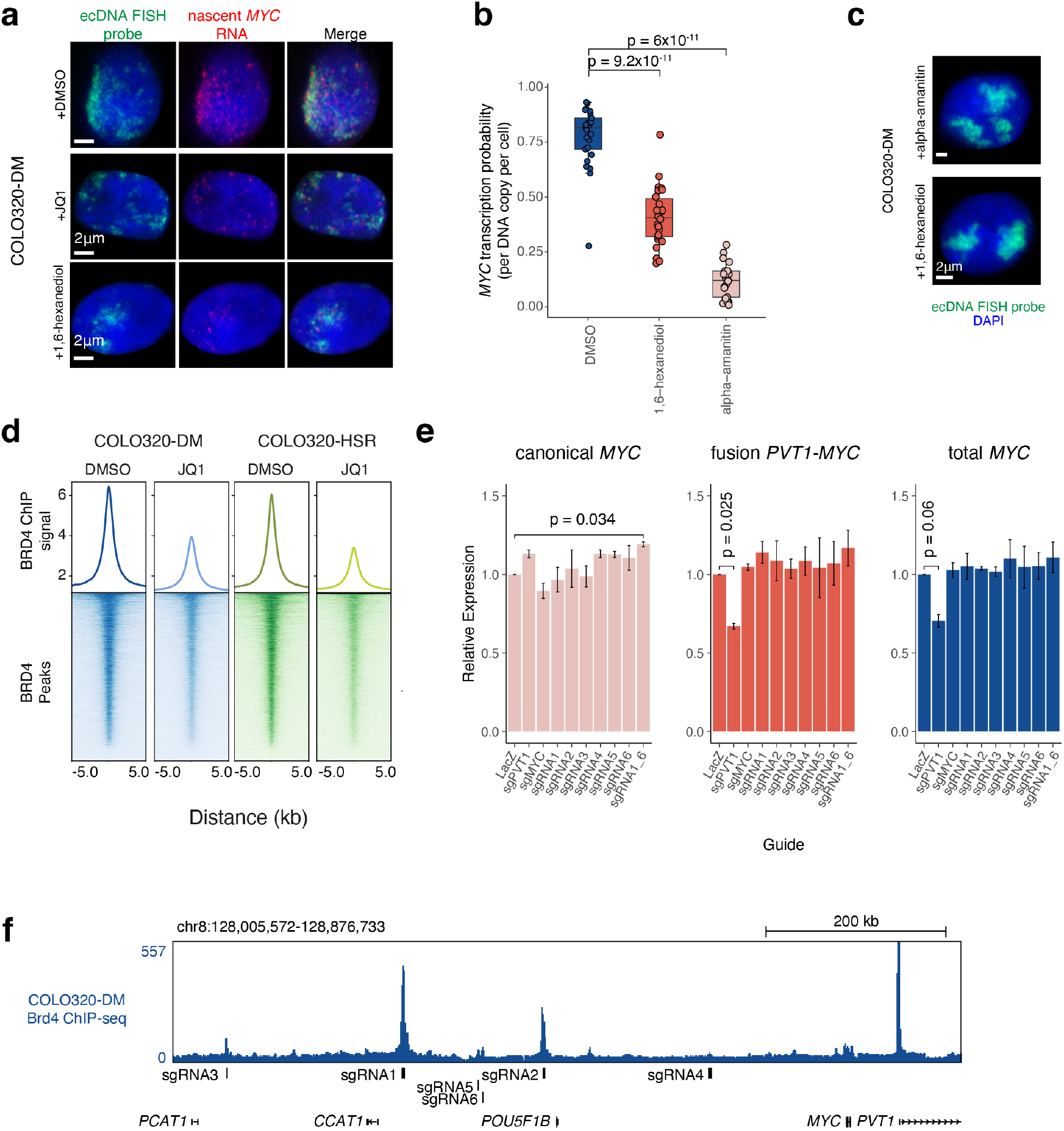
ecDNA transcription following 1,6-hexanediol treatment and inhibition of individual enhancers. **(a)** Representative image from combined DNA FISH for *MYC* ecDNA with nascent *MYC* RNA FISH in COLO320-DM cells treated with DMSO, 500 nM JQ1, or 1% 1,6-hexanediol for 6 hours. **(b)** Quantification of MYC transcription probability measured by nascent RNA FISH normalized to ecDNA copy number measured by DNA FISH for COLO320-DM cells treated either with DMSO, 1% 1,6-hexanediol, or 100 μg/mL alpha-amanitin for 6 hours. **(c)** Representative DNA FISH images for MYC ecDNA in interphase COLO320-DM cells treated with either 1% 1,6-hexanediol or 100 μg/mL alpha-amanitin for 6 hours. **(d)** Averaged BRD4 ChIP-seq signal (top) and heatmap of BRD4 ChIP-seq signal (bottom) over all BRD4 peaks for COLO320-DM and COLO320-HSR cells treated either with DMSO or 500 nM JQ1 for 6 hours. **(e)** Relative RNA expression measured by RT-qPCR for indicated transcripts in COLO320-DM cells stably expressing dCas9-KRAB and indicated sgRNAs. Canonical *MYC* was amplified with primers MYC_exon1_fw and MYC_exon2_rv; fusion PVT1-MYC was amplified with PVT1_exon1_fw and MYC_exon2_rv; total MYC was amplified with total_MYC_exon2_fw and total_MYC_exon2_rv. All primer sequences are in **Supplemental Table 1**. **(f)** BRD4 ChIP-seq coverage track at *MYC* locus in COLO320-DM cells with locations of sgRNA targeting sites of high BRD4 occupancy using for CRISPRi.

**Supplemental Figure 6.**
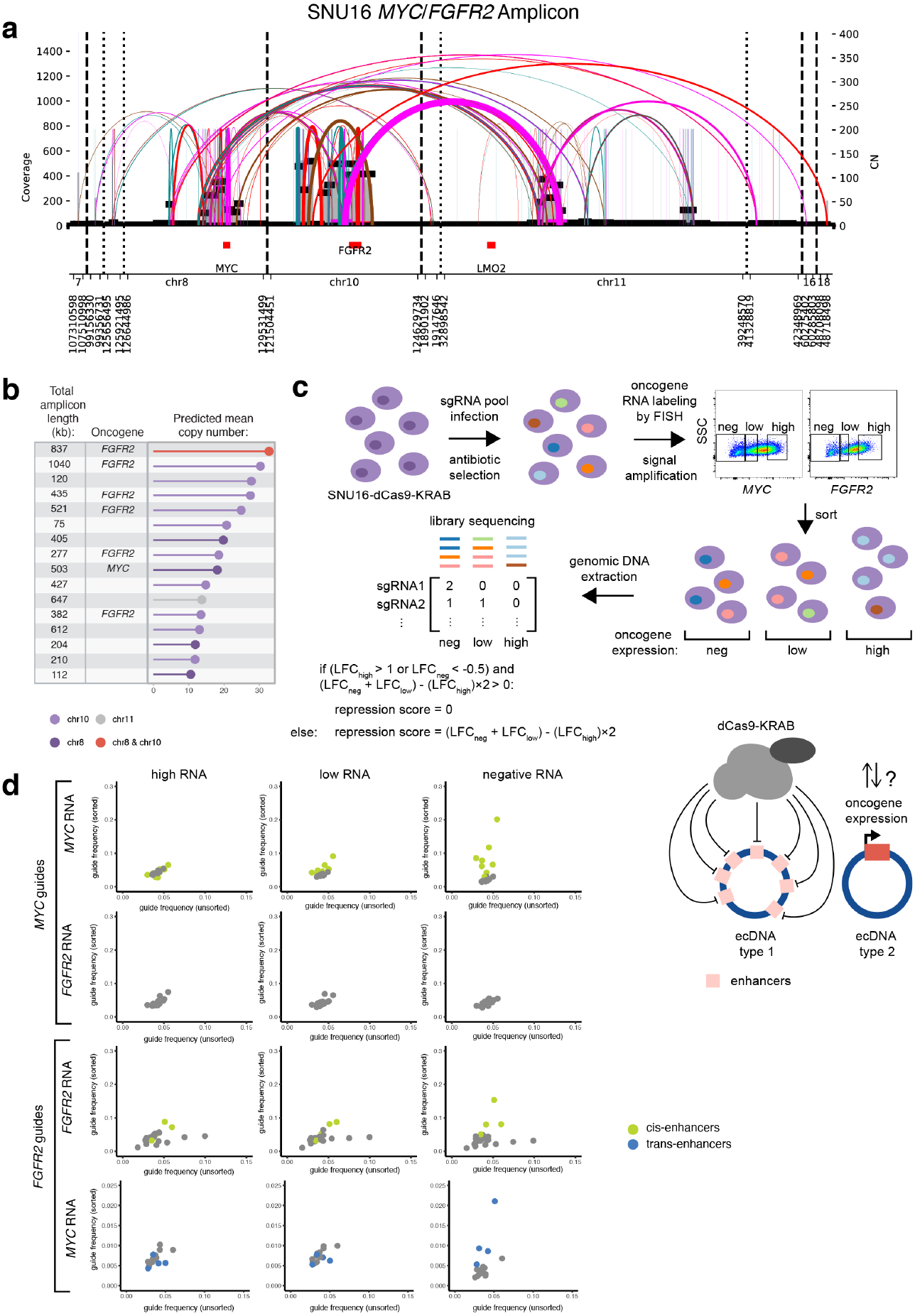
Structural rearrangements of dual oncogene ecDNAs and CRISPR interference of their putative enhancers. **(a)** Structural variant (SV) view of AmpliconArchitect (AA) reconstruction of the *MYC*/*FGFR2* amplicon in SNU16 cells. **(b)** Predicted amplicon length, oncogene content and mean copy number of *MYC*/*FGFR2* amplicon structures in SNU16 cells. Amplicon structures with over 50 kb in total length and over 10 predicted copies shown. **(c)** A schematic of CRISPR interference experiments perturbing potentially trans-acting enhancers in SNU16 cells. Single guide RNAs (sgRNAs) were designed to target putative trans-enhancers on *MYC*-bearing ecDNAs (ecDNA type 1) based on chromatin accessibility, and expression of *FGFR2* on distinct ecDNAs was measured (ecDNA type 2). A separate set of sgRNAs was designed to target putative trans-enhancers on *FGFR2*-bearing ecDNAs (ecDNA type 1), and expression of *MYC* was measured (ecDNA type 2) (bottom right). We performed these experiments in a pooled format. In brief, stable SNU16-dCas9-KRAB cells were generated and transduced with a lentiviral pool of sgRNAs targeting either the *MYC*-bearing ecDNAs or the *FGFR2*-bearing ecDNAs. Transduced cells were selected, oncogene RNA was labeled with FISH probes, FISH signals were amplified using PrimeFlow and assessed by flow cytometry. Cells with negative (neg), low or high RNA signals were sorted as different fractions, genomic DNA was extracted, and targeted libraries were prepared and sequenced. Relative abundances of sgRNAs were measured and a combined repression score was calculated by adding the log fold changes (LFCs) of each guide in the neg and low RNA fractions relative to unsorted cells, and subtracting from the LFC of each guide in the high RNA fraction. This repression score quantifies the degree to which each guide is enriched in the neg and low RNA fractions and depleted in the high RNA fraction. **(d)** Scatter plots showing relative frequencies of *MYC* ecDNA-targeting or *FGFR2* ecDNA-targeting guides in cells with high, low or negative levels of *MYC* or *FGFR2* RNA expression. Enhancers hits based on the combined repression scores are highlighted (cis in light green, trans in blue). We note that 8 of the *FGFR2* guides had consistently higher frequencies in *MYC*-sorted cells compared to unsorted cells. Because this enrichment is not differential between the high, low and negative RNA expressing cells, these guides do not fulfill our criteria outlined above for identifying guides with repressive effects on RNA expression and are therefore not categorized as hits. The y-axes in the bottom row was adjusted to exclude those guides.

**Supplemental Movie 1. Live cell imaging with untreated TetO-GFP COLO320-DM cells.** Snapshots of an untreated cell are shown over the course of 30 minutes. GFP labels TetO-knockin *MYC* ecDNAs.

**Supplemental Movie 2. Live cell imaging with DMSO-treated TetO-GFP COLO320-DM cells.** A control cell treated with DMSO was tracked over the course of 1 hour. GFP labels TetO-knockin *MYC* ecDNAs.

**Supplemental Movie 3. Live cell imaging with TetO-GFP COLO320-DM cells after JQ1 treatment.** A cell treated with 500 nM JQ1 was tracked over the course of 1 hour. GFP labels TetO-knockin *MYC* ecDNAs.

## METHODS

### Cell Culture

COLO320-DM, COLO320-HSR and HCC1569 cells were maintained in Roswell Park Memorial Institute 1640 (RPMI; Life Technologies, Cat# 11875-119) supplemented with 10% fetal bovine serum (FBS; Hyclone, Cat# SH30396.03) and 1% penicillin-streptomycin (pen-strep; Thermo Fisher, Cat# 15140-122). PC3 cells were maintained in Dulbecco’s Modified Eagle Medium (DMEM; Thermo Fisher, Cat# 11995073) supplemented with 10% FBS and 1% pen-strep. HK359 cells were maintained in DMEM/Nutrient Mixture F-12 (DMEM/F12 1:1; Gibco, Cat# 11320-082), B-27 Supplement (Gibco, Cat# 17504044), 1% pen-strep, GlutaMAX (Gibco, Cat# 35050061), human epidermal growth factor (EGF, 20 ng/ml; Sigma-Aldrich, E9644), human fibroblast growth factor (FGF, 20 ng/ml; Peprotech) and Heparin (5 ug/ml; Sigma-Aldrich, Cat# H3149-500KU). SNU16 cells were maintained in DMEM/F12 supplemented with 10% FBS and 1% pen-strep. All cells were cultured at 37°C with 5% CO_2_.

### RT-qPCR

RNA was extracted using RNeasy Plus mini Kit (QIAGEN 74136). Purified RNA was quantified by Nanodrop (Thermo Fisher). For RT-qPCR, 50 ng of RNA, 1X Brilliant II qRT-PCR mastermix with 1 uL RT/RNase block (Agilent 600825), and 200 nM forward and reverse primer were used. Each Ct value was measured using Lightcycler 480 (Roche) and each mean dCt was averaged from duplicate qRT-PCR reaction. Relative *MYC* RNA level (RT-qPCR primers MYC_exon3_fw and MYC_exon3_rv) was calculated by ddCt method compared to 18S and GAPDH controls (RT-qPCR primers GAPDH_fw, GAPDH_rv, 18S_fw, 18S_rv). The mean dCt value of each replicate was used to calculate p value using a Student’s t-test. Primer sequences are listed in **Supplemental Table 1**.

### Cell Viability Assays

Cells were plated in 96-well plates at 25,000 cells/well and incubated either with JQ1 (Sigma-Aldrich SML1524) at the indicated concentrations or an equivalent volume of DMSO for 48 hours. Cell viability was measured using the CellTiterGlo assay kit (Promega G7572) in triplicate with luminescence measured on SpectraMax M5 plate reader with an integration time of 1 second per well. Luminescence was normalized to the DMSO treated controls and p values calculated using a Student’s t-test.

### Lentivirus production

Lentiviruses were produced as previously described ^26^. Briefly, 4 million HEK293Ts per 10 cm plate were plated the evening before transfection. Helper plasmids, pMD2.G and psPAX2, were transfected along with the vector plasmid using Lipofectamine 3000 (Thermo Fisher, Cat# L3000) according to the manufacturer’s instructions. Supernatants containing lentivirus were harvested 48 hours later, filtered with a 0.45 um filter and concentrated using Lenti-X concentrator (Clontech, Cat#631232) and stored at 80°C.

### Stable CRISPR cell line generation

The pHR-SFFV-dCas9-BFP-KRAB (Addgene, Cat# 46911) plasmid was modified to dCas9-BFP-KRAB-2A-Blast as previously described ^26^. Lentivirus was produced using the modified vector plasmid. Cells were transduced with lentivirus, incubated for 2 days, selected with 1ug/ml blasticidin for 10-14 days, and BFP expression was analyzed by flow cytometry. To generate stable, monoclonal dCas9-KRAB cell lines, single BFP-positive cell clones were sorted into 96-well plates and expanded. Vector expression was validated by flow cytometry.

### CRISPR interference

sgRNAs were designed using the Broad Institute sgRNA designer online tool (https://portals.broadinstitute.org/gpp/public/analysis-tools/sgrna-design). An additional guanine was appended to each of the protospacers that do not start with a guanine. sgRNAs were cloned into either mU6(modified)-sgRNA-Puromycin-mCherry or mU6(modified)-sgRNA-Puromycin-EGFP previously generated ^26^ and lentiviruses were produced. To evaluate the effects of CRISPR interference on gene expression, cells were transduced with sgRNA lentiviruses, incubated for 2 days, selected with 0.5ug/ml puromycin for 4 days, and BFP, GFP and/or mCherry expressions were assessed by flow cytometry. Cells were harvested for RT-qPCR assays.

For the pooled experiments in SNU16, sgRNAs targeting bulk ATAC-seq peaks were designed, cloned, pooled and lentiviruses were produced. SNU16-dCas9-KRAB cells were transduced with the lentiviral guide pool, incubated for 2 days, selected with puromycin for 4 days, and RNA FISH flow was performed for *MYC* and *FGFR2* using the PrimeFlow™ RNA Assay Kit (Thermo Fisher) following the manufacturer’s protocol and corresponding probe sets (*MYC*: VA1-6000107-PF; *FGFR2*: VA1-14785-PF). Cells were sorted by fluorescence-activated cell sorting (FACS), and genomic DNA was extracted as previously described ^64^. Libraries were prepared using 3 rounds of PCR. The first round was performed using primers sgRNA_backbone_outer_fw and sgRNA_backbone_outer_rv to amplify guide sequences. The second round was a nested PCR with primers p5_mU6_0nt_stagger, p5_mU6_1nt_stagger, p5_mU6_2nt_stagger, p5_mU6_3nt_stagger mixed at equimolar ratios and reverse primer p7adpt_spRNAl105nt_rev. Finally, sequencing indices were attached in the third PCR using primers that anneal to the adaptors and contain Illumina Truseq dual index primers. Initial amplification and nested PCR primer sequences are listed in **Supplemental Table 1**. Amplified product sizes were validated on a gel, and the products were purified using SPRIselect reagent kit (Beckman Coulter, Cat# B23318) at 1.2x sample volumes following the manufacturer’s protocol. Libraries were sequenced on an Illumina Miseq with paired-end 75 bp read lengths.

Relative abundances of sgRNAs were measured and compared using MAGeCK ^65^. sgRNA counts were obtained using the “mageck count” command, and differential enrichment analysis was performed for each sorted cell fraction compared to unsorted cells using the “mageck test” command. Log fold changes (LFCs) relative to unsorted cells were obtained for each guide, and a combined repression score was calculated as (LFC_neg_ + LFC_low_) – (LFC_high_) × 2. In cases where LFChigh > 1 or LFCneg < −0.5 and the repression scores were above zero, we adjusted the repression scores to zero as the guides were considered to be enriched in cells with high expression and/or depleted in cells with low expression, i.e. they were non-repressive. All sgRNA sequences are listed in **Supplemental Table 2**.

### Metaphase chromosome spread

Cells in metaphase were prepared by KaryoMAX (Gibco) treatment at 0.1 ug/ml for 3 hr. Single-cell suspension was then collected and washed by PBS, and treated with 75 mM KCl for 15-30 min. Samples were then fixed by 3:1 methanol:glacial acetic acid, v/v and washed for an additional three times with the fixative. Finally, the cell pellet resuspended in the fixative was dropped onto a humidified slide.

### DNA FISH

Slides containing fixed cells in interphase or metaphase were briefly equilibrated by 2X SSC, followed by dehydration in 70%, 85%, and 100% ethanol for 2 min each. FISH probes in hybridization buffer (Empire Genomics) were added onto the slide, and the sample was covered by a coverslip then denatured at 75°C for 1 min on a hotplate, and hybridized at 37°C overnight. The coverslip was then removed, and the sample was washed one time by 0.4X SSC with 0.3% IGEPAL, and two times by 2X SSC with 0.1% IGEPAL, for 2 min each. DNA was stained with DAPI and washed with 2X SSC. Finally, the sample was mounted by mounting media (Molecular Probes) before imaging.

The Oligopaint FISH probe libraries were constructed as described previously ^66^. Each oligo consists of a 40 nucleotide (nt) homology to the hg19 genome assemble designed from the algorithm developed from the laboratory of Dr.Ting Wu (https://oligopaints.hms.harvard.edu/). Each library subpool consists of a unique sets of primer pairs for orthogonal PCR amplification and a 20 nt T7 promoter sequence for *in vitro* transcription and a 20 nt region for reverse transcription. Individual Oligopaint probes were generated by PCR amplification, *in vitro* transcription, and reverse transcription, in which ssDNA oligos conjugated with ATTO488 and ATTO647 fluorophores were introduced during the reverse transcription step. The Oligopaint covered genomic regions (hg19) used in this study are as follows: chr8:116967673-118566852 (hg19_COLO_nonecDNA_1.5Mbp), chr8:127435083-129017969 (hg19_COLO_ecDNA_1.5Mbp), chr8:128729248-128831223 (hg19_PC3_ecDNA1_100kb). A ssDNA oligo pool was ordered and synthesized from Twist Bioscience (San Francisco, CA). 15mm #1.5 round glass coverslips (Electron Microscopy Sciences) were pre-rinsed with anhydrous ethanol for 5min, air dried, and coated with L-poly lysine solution (100ug/mL) for at least 2 hours. Fully dissociated ColoDM320 or PC3 cells were seeded onto the coverslips and recovered for at least 6 hours before experiments. Cells were fixed with 4% (v/v) methanol free paraformaldehyde diluted in 1X PBS at room temperature for 10min. Then cells were washed 2X with 1XPBS and permeabilized in 0.5% Triton-X100 in 1XPBS for 30min. After 2X wash in 1XPBS, cells were treated with 0.1M HCl for 5min, followed by 3X washes with 2XSSC and 30 min incubation in 2X SSC + 0.1% Tween20 (2XSSCT) + 50% (v/v) formamide (EMD Millipore, cat#S4117). For each sample, we prepare 25ul hybridization mixture containing 2XSSCT+ 50% formamide +10% Dextran sulfate (EMD Millipore, cat#S4030) supplemented with 0.5μl 10mg/mL RNaseA (Thermo Fisher Scientific, cat# 12091-021) +0.5μl 10mg/mL salmon sperm DNA (Thermo Fisher Scientific, cat# 15632011) and 20pmol probes with distinct fluorophores. The probe mixture was thoroughly mixed by vortexing, and briefly microcentrifuged. The hybridization mix was transferred directly onto the coverslip which was inverted facing a clean slide. The coverslip was sealed onto the slide by adding a layer of rubber cement around the edges. Each slide was denatured at 78°C for 4 min followed by transferring to a humidified hybridization chamber and incubated at 42°C for 16 hours in a heated incubator. After hybridization, samples were washed 2X for 15 minutes in pre-warmed 2XSSCT at 60 °C and then were further incubated at 2XSSCT for 10min at RT, at 0.2XSSC for 10min at RT, at 1XPBS for 2X5min with DNA counterstaining with DAPI. Then coverslips were mounted on slides with Prolong Diamond Antifade Mountant (Thermo Fisher Scientific Cat#P36961) for imaging acquisition.

### Nascent RNA FISH

To quantify the MYC gene expression on the ecDNAs, we ordered the RNA FISH probes conjugated with a Quasar 570 dye (Biosearch Technologies) targeting to the intronic region of human (hg19) MYC gene for detection of nascent RNA transcript. We also ordered the RNA FISH probes conjugated with a Quasar 670 dye targeting to the exonic region of human MYC gene for detection of both mature and nascent RNA transcripts. For simultaneous detection of both ecDNA and MYC transcription, 125nM RNA FISH probes was mixed with the DNA FISH probes (100kb probe instead of the 1.5Mbp probe) together in the hybridization buffer without RNaseA and incubated at 37°C overnight for ~16 hours. After hybridization, samples were washed 2X for 15 minutes in pre-warmed 2XSSCT at 37 °C and then were further incubated at 2XSSCT for 10min at RT, at 0.2XSSC for 10min at RT, at 1XPBS for 2X5min with DNA counterstaining with DAPI. Then coverslips were mounted on slides with Prolong Diamond Antifade Mountant for imaging acquisition.

### Microscopy

DNA FISH images were acquired either with conventional fluorescence microscopy or confocal microscopy. Conventional fluorescence microscopy was performed using an Olympus BX43 microscope, and images were acquired with a QiClick cooled camera. Confocal microscopy was performed using a Leica SP8 microscope with lightning deconvolution (UCSD School of Medicine Microscopy Core). Z-stacks were acquired over an average depth of approximately 8μm, with roughly 0.6μm step size.

DNA/RNA FISH images were acquired on the ZEISS LSM 880 Inverted Confocal microscope attached with an Airyscan 32 GaAsP PMT area detector. Before imaging, the beam position was calibrated centering on the 32 detector array. Images were taken under the Airyscan SR mode with a Plan Apochromat 63X/NA1.40 oil objective in a lens immersion medium having a refractive index 1.515 at 30oC. We used 405nm (Excitation wavelength) and 460nm (Emission wavelength) for the DAPI channel, 488nm (Excitation wavelength) and 525nm (Emission wavelength) for the ATTO488 channel, 561nm (Excitation wavelength) and 579nm (Emission wavelength) for the Quasar570 channel and 633nm (Excitation wavelength) and 654nm (Emission wavelength) for the ATTO647 channel. Z-stacks were acquired with the optimal z sectioning thickness ~200nm, followed by post-processing using the provided algorithm from ZEISS LSM880 platform.

### Generation of ecDNA-TetO array for live cell imaging

sgRNA was designed by E-CRISP (http://www.e-crisp.org/E-CRISP/designcrispr.html) targeting ~0.5kb upstream of *MYC* transcription start site. The sgRNA sequence is listed in **Supplemental Table 2**. The sgRNA was cloned into the modified pX330 (Addgene, Cat# 42230) construct co-expressing wild type SpCas9 and a PGK-Venus cassette. ~500bp homology arms were PCR amplified from COLO320-DM cells and cloned into a pUC19 donor vector together with ~100 copies of TetO array and a blasticidin selection cassette. 2 ug of the donor vector and 1 ug of the sgRNA vector were transfected into COLO320-DM cells by lipofectamine 3000 and blasticidin (10 ug/ml) selection was applied after 7 days. Individual clones were selected, genotyped by PCR and verified by Sanger sequencing before being tested for imaging. To detect TetO array-labeled ecDNA molecules, we transiently expressed TetR-eGFP as previously reported ^67^ and performed imaging experiments two days after transfection.

### Image Analysis

To analyze the clustering of ecDNAs, we applied the autocorrelation function as described previously ^68^ in Matlab (2019). Specifically, the pair auto-correlation function 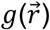 was calculated by the fast Fourier transform (FFT) method described by the equations below.

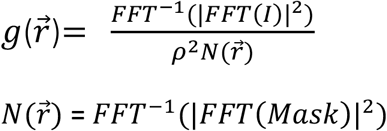

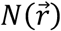 is the auto-correlation of a mask matrix that has the value of 1 inside the nucleus used for normalization. The fast Fourier transform and its inverse (*FFT* and *FFT*^−1^) were computed by fft2() and ifft2() functions in Matlab, respectively. Autocorrelation functions were calculated first by converting the Cartesian coordinates to polar coordinates by Matlab cart2pol() function, binning by radius and by averaging within the assigned bins. For comparing auto-correlation with transcription probability, the value of the auto-correlation function at radius of 0 pixels (g(0)) was used to represent the degree of spatial clustering. The g(0) values were also used for calculating statistical significance among groups.

To characterize the ecDNA shape and size, we employed the synthetic model—Surfaces object from Imaris and applied a Gaussian filter (σ = 1 voxel in xy) before the downstream segmentation and quantification. To measure the number of ecDNA or nascent transcripts, we localized the voxels corresponding to the local maximum of identified DNA or RNA FISH signal using the Imaris spots function module.

Colocalization analysis was performed using confocal images of both metaphase and interphase nuclei from the same slides. Images were split into the two FISH colors, and background fluorescence was removed manually for each channel. Colocalization for each nucleus was quantified using the ImageJ-Colocalization Threshold program. Analysis was performed across all z-stacks for each nucleus. Manders coefficient (fraction of *MYC* signal colocalized compared to total MYC signal) was used to quantify colocalization.

### Whole Genome Sequencing

Whole genome sequencing data from COLO320-DM, COLO320-HSR and PC3 cells were generated by a previously published study ^1^ and obtained from the NCBI Sequence Read Archive, under BioProject accession PRJNA506071. Whole genome sequencing data from SNU16 cells was generated by a previously published study ^69^ and obtained from the NCBI Sequence Read Archive, under BioProject accession PRJNA523380. Whole genome sequencing data from HK359 cells was generated by a previously published study ^6^ and obtained from the NCBI Sequence Read Archive, under BioProject accession PRJNA338012.

### Long Read Sequencing

Genomic DNA from COLO320-DM cells was extracted using a MagAttract HMW DNA Kit (Qiagen 67563) and prepared for long read sequencing using a Ligation Sequencing Kit (Oxford Nanopore Technologies SQK-LSK109) according to the manufacturer’s instructions. Sequencing was performed on a MinION (Oxford Nanopore Technologies).

### RNA-seq Library Preparation

COLO320-DM cells were transfected with Alt-R^®^ S.p. Cas9 Nuclease V3 (IDT, Cat# 1081058) complexed with a non-targeting control sgRNA (Synthego) with a LacZ sequence following Synthego’s RNP transfection protocol using the Neon Transfection System (ThermoFisher, Cat# MPK5000). 500,000 to 1 million cells were harvested, and RNA was extracted using RNeasy Plus mini Kit (QIAGEN 74136). Genomic DNA was removed from samples using the TURBO DNA-free kit (ThermoFisher, Cat# AM1907), and RNA-seq libraries were prepared using the TruSeq Stranded mRNA Library Prep (Illumina, Cat# 20020595) following the manufacturer’s protocol. RNA-seq libraries were sequenced on an Illumina HiSeq 4000 with paired-end 75 bp read lengths.

### ChIP-seq Library Preparation

Three million cells per replicate were fixed in 1% formaldehyde for 10 minutes at room temperature with rotation and then quenched with 0.125 M glycine for 10 minutes at room temperature with rotation. Fixed cells were pelleted at 800xg for 5 minutes at 4°C and washed twice with cold PBS before storing at −80°C. Pellets were thawed and membrane lysis performed in 5 mL LB1 (50 mM HEPES pH 8.0, 140 mM NaCl, 1 mM EDTA, 10% glycerol, 0.5% NP-40, 0.25% Triton X-100, 1 mM PMSF, Roche protease inhibitors 11836170001) for 10 min at 4°C with rotation. Nuclei were pelleted at 1350xg for 5 min at 4°C and lysed in 5 mL LB2 (10 mM Tris-Cl pH 8.0, 5 M, 200 mM NaCl, 1 mM EDTA, 0.5 mM EGTA, 1 mM PMSF, Roche protease inhibitors) for 10 min at RT with rotation. Chromatin was pelleted at 1350xg for 5 min at 4°C and resuspended in 1 mL of TE Buffer + 0.1% SDS before sonication on a Covaris E220. Samples were clarified by spinning at 16,000xg for 10 min at 4°C. Supernatant was transferred to a new tube and diluted with 1 volume of IP Dilution Buffer (10 mM Tris pH 8.0, 1 mM EDTA, 200 mM NaCl, 1 mM EGTA. 0.2% Na-DOC, 1% Na-Laurylsarcosine, 2% Triton X-100). Following addition of 20 ng spike-in chromatin (Active Motif 61686) and 2 μg spike-in antibody (Active Motif 53083), 50 μL of sheared chromatin was reserved as input and ChIP performed overnight at 4°C with rotation with 7.5 μg of antibody per IP: H3K27Ac (Abcam ab4729), BRD4 (Bethyl Laboratories A301-985A100).

100 μL Protein G Dynabeads per ChIP were washed 3X in 0.5% BSA in PBS and then bound to antibody bound chromatin for 4 hours at 4°C with rotation. Antibody bound chromatin was washed on a magnet 5X with RIPA Wash Buffer (50 mM HEPES pH 8.0, 500 mM LiCl, 1 mM EDTA, 1% NP-40, 0.7% Na-Deoxycholate) and once with 1 mL TE Buffer (10 mM Tris-Cl pH 8.0, 1 mM EDTA) with 500 mM NaCl. Washed beads were resuspended in 200 mL ChIP Elution Buffer (50 mM Tris-Cl pH 8.0, 10 mM EDTA, 1% SDS) and chromatin was eluted following incubation at 65°C for 15 min. Supernatant and input chromatin were removed to fresh tubes and reverse cross-linked at 65°C overnight. Samples were diluted with 200 mL TE Buffer, treated with 0.2 mg/mL RNase A (QIAGEN 19101) for 2 hours at 37°C, then 0.2 mg/mL Proteinase K (New England Biolabs P8107S) for 30 min at 55°C. DNA was purified using the ChIP DNA Clean & Concentrator kit (Zymo Research D5205). ChIP sequencing libraries were prepared using the NEBNext Ultra II DNA Library Prep Kit for Illumina (New England Biolabs E7645S) with dual indexing (New England Biolabs E7600S) following the manufacturer’s instructions. ChIP-seq libraries were sequenced on an Illumina HiSeq 4000 with paired-end 76 bp read lengths.

### HiChIP Library Preparation

One to four million cells were fixed in 1% formaldehyde in aliquots of one million cells each for 10 minutes at room temperature. HiChIP was performed as previously described ^40^ using antibodies against H3K27ac (Abcam ab4729) with the following optimizations ^70^: SDS treatment at 62°C for 5 min; restriction digest with MboI for 15 min; instead of heat inactivation of MboI restriction enzyme, nuclei were washed twice with 1X restriction enzyme buffer; biotin fill-in reaction incubation at 37°C for 15 minutes; ligation at room temperature for 2 hours. HiChIP libraries were sequenced on an Illumina HiSeq 4000 with paired-end 76 bp read lengths.

### Single-Cell Paired RNA and ATAC-seq Library Preparation

Single-cell paired RNA and ATAC-seq libraries for COLO320-DM and COLO320-HSR were generated on the 10x Chromium Single-Cell Multiome ATAC + Gene Expression platform following the manufacturer’s protocol and sequenced on an Illumina NovaSeq 6000.

### Long Read Sequencing Data Processing

Bases were called from fast5 files using guppy (Oxford Nanopore Technologies, version 2.3.7). Reads were then aligned using NGMLR ^71^ (version 0.2.7) with the following parameters: -x ont --no-lowqualitysplit. Structural variants were called using Sniffles ^71^ (version 1.0.11) using the following parameters: -s 1 --report_BND --report_seq. Read information regarding mapping locations and junctions present in individual reads stored in QS and QE read tags was extracted using samtools and visualized in R.

### RNA-seq Data Processing

Paired-end reads were aligned to the hg19 genome using STAR-Fusion ^72^ (version 1.6.0) and the genome build GRCh37_gencode_v19_CTAT_lib_Mar272019.plug-n-play. Number of reads supporting the *PVT1-MYC* fusion transcript were obtained from the “star-fusion.fusion_predictions.abridged.tsv” output file and the junction read counts and spanning fragment counts were combined. Reads supporting the canonical *MYC* exon 1-2 junction were obtained using the Gviz package in R ^73^ in a sashimi plot.

### ChIP-seq Data Processing

Paired-end reads were aligned to the hg19 genome using Bowtie2 ^74^ (version 2.3.4.1) with the --very-sensitive option following adapter trimming with Trimmomatic ^75^ (version 0.39). Reads with MAPQ values less than 10 were filtered using samtools and PCR duplicates removed using Picard’s MarkDuplicates. MACS2 ^76^ (version 2.1.1.20160309) was used for peak calling with the following parameters: macs2 callpeak -t chip_bed -c input_bed -n output_file -f BED -g hs -q 0.01 --nomodel --shift 0. A reproducible peak set across biological replicates was defined using the IDR framework (version 2.0.4.2). Reproducible peaks from all samples were then merged to create a union peak set. ChIP-seq signal was converted to bigwig format for visualization using deepTools bamCoverage ^77^ (version 3.3.1) with the following parameters: --bs 5 --smoothLength 105 --normalizeUsing CPM --scaleFactor 10. Enrichment of ChIP signal at peaks was performed using deepTools computeMatrix.

### HiChIP Data Processing

HiChIP data were processed as described previously ^40^. Briefly, paired end reads were aligned to the hg19 genome using the HiC-Pro pipeline (version 2.11.0) ^78^. Default settings were used to remove duplicate reads, assign reads to MboI restriction fragments, filter for valid interactions, and generate binned interaction matrices. The Juicer pipeline’s HiCCUPS tool and FitHiChIP were used to identify loops ^79,80^. Filtered read pairs from the HiC-Pro pipeline were converted into .hic format files and input into HiCCUPS using default settings. Dangling end, self-circularized, and re-ligation read pairs were merged with valid read pairs to create a 1D signal bed file. FitHiChIP was used to identify “peak-to-all” interactions at 10 kb resolution using peaks called from the one-dimensional HiChIP data. A lower distance threshold of 20 kb was used. Bias correction was performed using coverage specific bias. HiChIP contact matrices stored in .hic files were visualized in Juicebox using square root coverage normalization. Virtual 4C plots were generated from dumped matrices generated with Juicebox. The Juicebox tools dump command was used to extract the chromosome of interest from the .hic file. The interaction profile of a 10-kb bin containing the anchor was then plotted in R following normalization by total number of valid read pairs and anchor bin signal.

### Single-Cell Paired RNA and ATAC-seq Data Processing

A custom reference package for hg19 was created using cellranger-arc mkref (10x Genomics, version 1.0.0). The single-cell paired RNA and ATAC-seq reads were aligned to the hg19 reference genome using cellranger-arc count (10x Genomics, version 1.0.0).

### Combined single-cell RNA and ATAC-seq analysis

Subsequent analyses on RNA were performed using Seurat ^81^, and those on ATAC-seq were performed using ArchR ^31^. Cells with more than 200 unique RNA features, less than 20% mitochondrial RNA reads, less than 50,000 total RNA reads were retained for further analyses. Doublets were removed using ArchR.

Raw RNA counts were normalized using the NormalizeData function, scaled using the ScaleData function, and the data were visualized on a UMAP using the first 30 principal components. Dimensionality reduction for the ATAC-seq data were performed using Iterative Latent Semantic Indexing (LSI) with the addIterativeLSI function in ArchR. To impute accessibility gene scores, we used addImputeWeights to add impute weights and plotEmbedding to visualize scores. To compare the accessibility gene scores for *MYC* with *MYC* RNA expression, getMatrixFromProject was used to extract the gene score matrix and the normalized RNA data were used.

To identify variable ATAC-seq peaks on COLO320-DM and COLO320-HSR amplicons, we first calculated amplicon copy numbers based on background ATAC-seq signals as previously described, using a sliding window of five megabases moving in one-megabase increments across the reference genome ^82^. We used the copy number z scores calculated for the chr8:124000001-129000000 interval for estimating copy numbers of *MYC*-bearing ecDNAs in COLO320-DM and *MYC*-bearing chromosomal HSRs in COLO320-HSR. We then incorporated these estimated copy numbers into the variable peak analysis as follows. COLO320-DM and COLO320-HSR cells were separately assigned into 20 bins based on their RNA expression of *MYC*. Next, pseudo-bulk replicates for ATAC-seq data were created using the addGroupCoverages function grouped by MYC RNA quantile bins. ATAC-seq peaks were called using addReproduciblePeakSet for each quantile bin, and peak matrices were added using addPeakMatrix. Differential peak testing was performed between the top and the bottom RNA quantile bins using getMarkerFeatures. A false discovery rate cutoff of 1e-15 was imposed. The mean copy number z score for each quantile bin was then calculated and a copy number fold change between the top and bottom bin was computed. Finally, we filtered on significantly differential peaks that are located in chr8:127432631-129010071 and have fold changes above the calculated copy number fold change multiplied by 1.5.

## Data Availability

ChIP-seq, HiChIP and single cell multiome ATAC + gene expression data generated in this study have been deposited in GEO and are available under accession number GSE159986. GEO dataset will be made publicly available upon publication of the peer-reviewed paper.

## Code Availability

All custom code used in this work is available from the corresponding authors upon reasonable request.

## Acknowledgements

We thank members of the Chang, Liu, Mischel, and Bafna laboratories for discussions and X. Ji, D. Wagh and J. Coller at the Stanford Functional Genomics Facility. H.Y.C. was supported by NIH R35-CA209919 and RM1-HG007735. K.L.H. was supported by the Stanford Graduate Fellowship. K.E.Y. was supported by the National Science Foundation Graduate Research Fellowship Program (NSF DGE-1656518), a Stanford Graduate Fellowship, and a NCI Predoctoral to Postdoctoral Fellow Transition Award (NIH F99CA253729). Cell sorting for this project was done on instruments in the Stanford Shared FACS Facility. Sequencing was performed by the Stanford Functional Genomics Facility (supported by NIH grants S10OD018220 and 1S10OD021763). Microscopy was performed on instruments in the UCSD Microscopy Core (supported by NINDS NS047101). Z.L. is a Janelia Group Leader and H.Y.C. and R.T. are Investigators of the Howard Hughes Medical Institute.

## Author Contributions

K.L.H., K.E.Y., and H.Y.C. conceived the project. K.L.H., K.E.Y., L.X., S.W., J.T.L., C.V.D., K.K., J.T., and R.L. performed experiments. K.L.H., K.E.Y., L.X., S.W., J.T.L. analyzed data with input from M.R.C., J.M.G., and U.R. Q.S., J.C.R., R.T., V.B., P.S.M., Z.L., and H.Y.C. guided data analysis and provided feedback on experimental design. K.L.H., K.E.Y., and H.Y.C. wrote the manuscript with input from all authors.

## Materials & Correspondence

Correspondence and requests for materials should be addressed to Howard Y. Chang.

## Supplemental Tables

**Supplemental Table 1.**
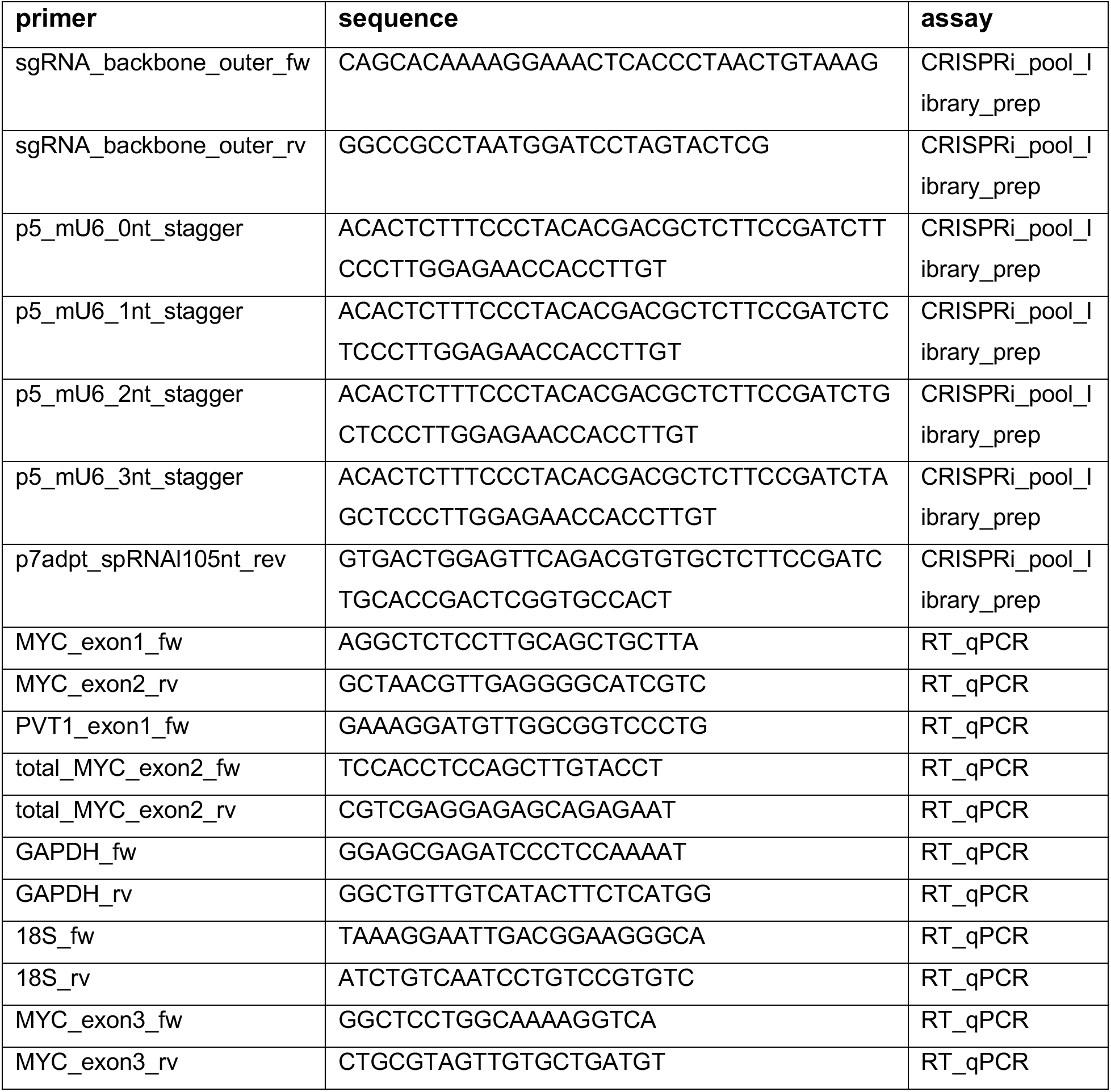
Primer sequences. All primers used for library amplification or RT-qPCR are listed.

**Supplemental Table 2.**
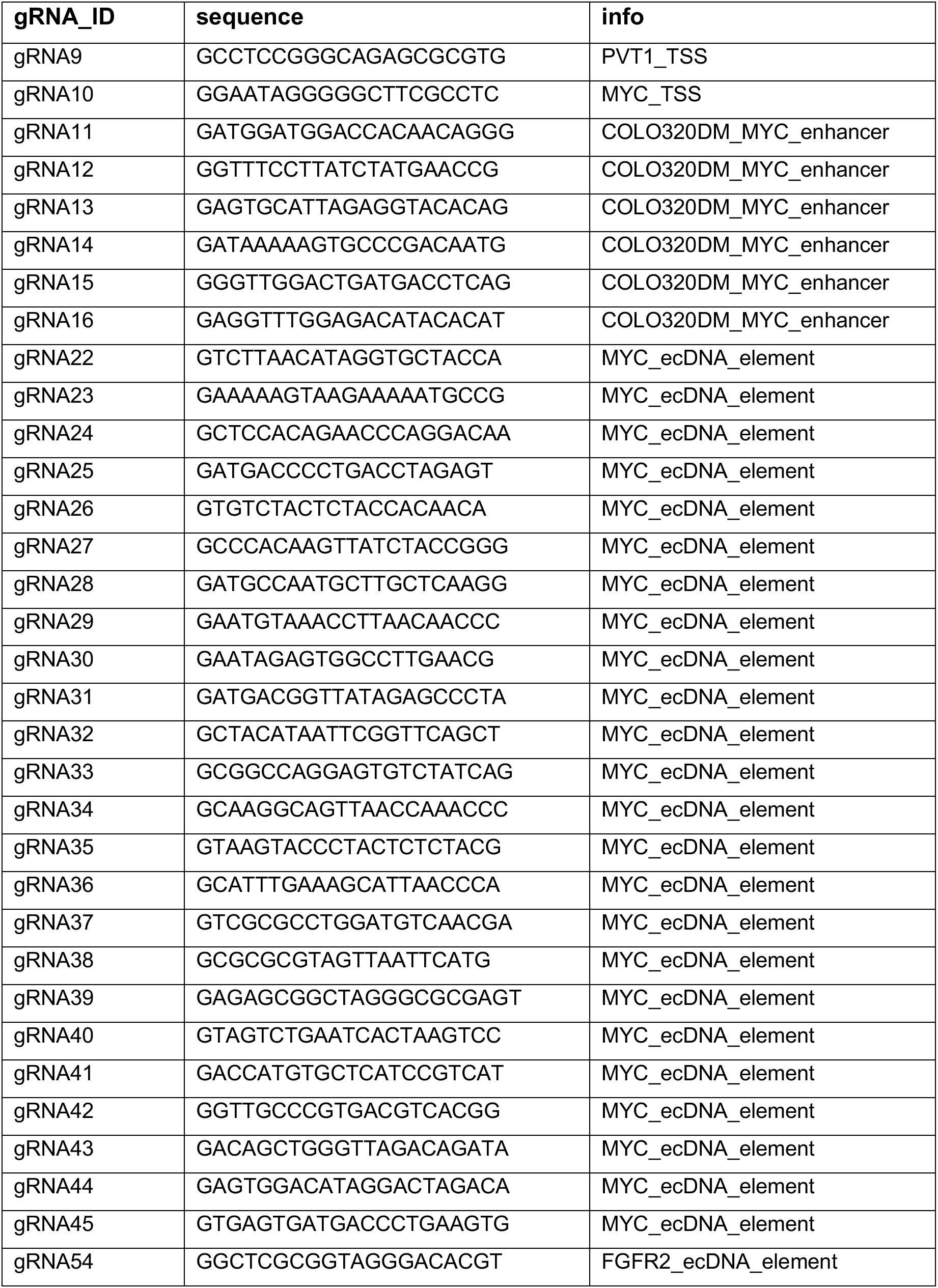
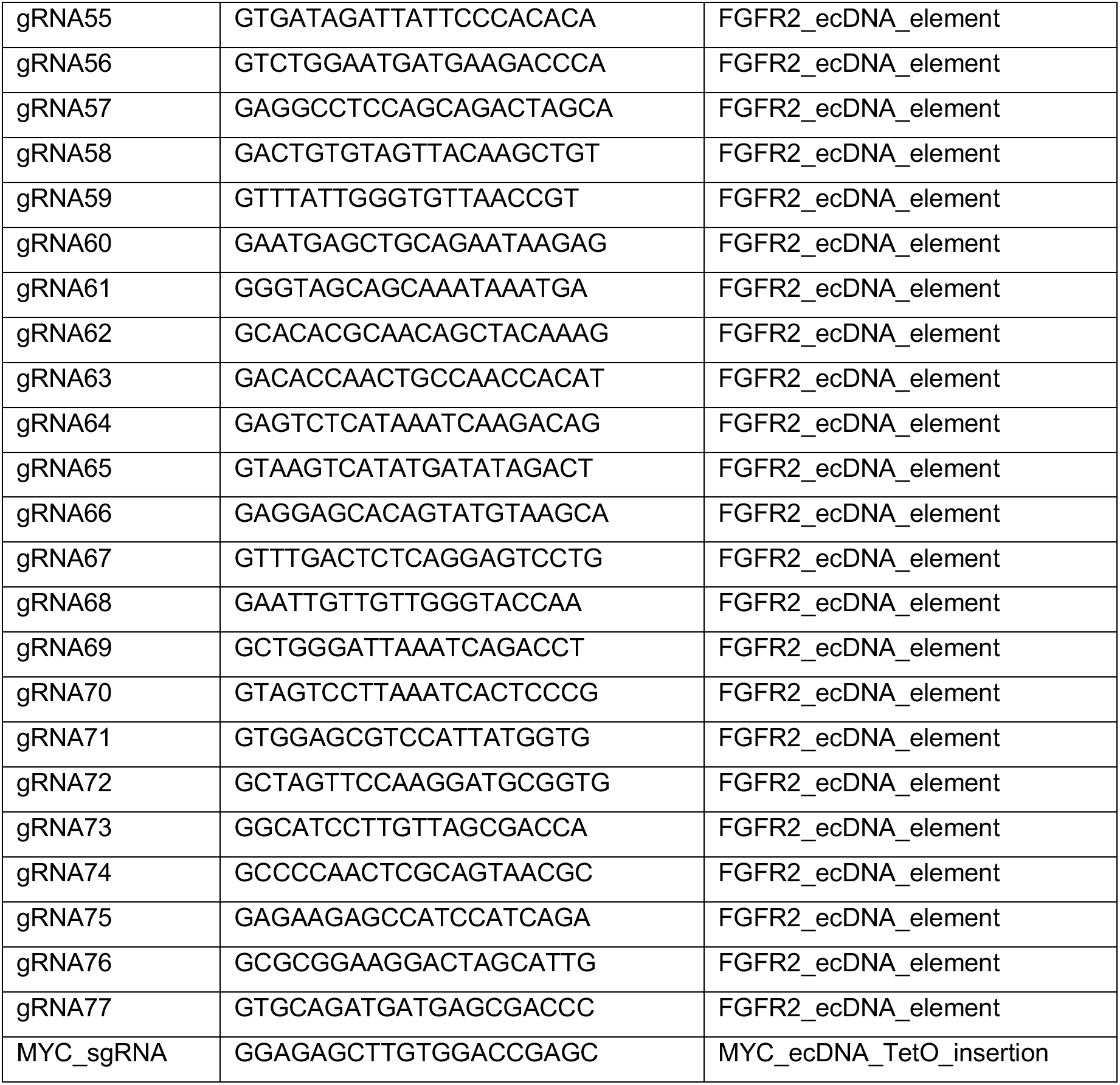
sgRNA sequences. All sgRNAs used in the CRISPR interference studies and ecDNA editing for TetO insertion are listed.

